# Learning produces a hippocampal cognitive map in the form of an orthogonalized state machine

**DOI:** 10.1101/2023.08.03.551900

**Authors:** Weinan Sun, Johan Winnubst, Maanasa Natrajan, Chongxi Lai, Koichiro Kajikawa, Michalis Michaelos, Rachel Gattoni, Carsen Stringer, Daniel Flickinger, James E. Fitzgerald, Nelson Spruston

## Abstract

Cognitive maps confer animals with flexible intelligence by representing spatial, temporal, and abstract relationships that can be used to shape thought, planning, and behavior. Cognitive maps have been observed in the hippocampus, but their algorithmic form and the processes by which they are learned remain obscure. Here, we employed large-scale, longitudinal two-photon calcium imaging to record activity from thousands of neurons in the CA1 region of the hippocampus while mice learned to efficiently collect rewards from two subtly different versions of linear tracks in virtual reality. The results provide a detailed view of the formation of a cognitive map in the hippocampus. Throughout learning, both the animal behavior and hippocampal neural activity progressed through multiple intermediate stages, gradually revealing improved task representation that mirrored improved behavioral efficiency. The learning process led to progressive decorrelations in initially similar hippocampal neural activity within and across tracks, ultimately resulting in orthogonalized representations resembling a state machine capturing the inherent structure of the task. We show that a Hidden Markov Model (HMM) and a biologically plausible recurrent neural network trained using Hebbian learning can both capture core aspects of the learning dynamics and the orthogonalized representational structure in neural activity. In contrast, we show that gradient-based learning of sequence models such as Long Short-Term Memory networks (LSTMs) and Transformers do not naturally produce such orthogonalized representations. We further demonstrate that mice exhibited adaptive behavior in novel task settings, with neural activity reflecting flexible deployment of the state machine. These findings shed light on the mathematical form of cognitive maps, the learning rules that sculpt them, and the algorithms that promote adaptive behavior in animals. The work thus charts a course toward a deeper understanding of biological intelligence and offers insights toward developing more robust learning algorithms in artificial intelligence.

## INTRODUCTION

Intelligence, at its core, manifests in an organism’s or an agent’s ability to engage dynamically with its environment, interpret information, adjust to unfamiliar situations, and execute complex tasks. A central concept in the study of natural and artificial intelligence (AI) is the notion of ‘internal models’. These models convert external world observations into a well-organized representation, thereby enabling adaptive behavior. In neuroscience, a notable example of internal models is the concept of ‘cognitive maps’. Conceptualized early in the twentieth century, cognitive maps are believed to be neural constructs that enable animals to comprehend their environment and understand the interactions between their bodies and the external world, which supports efficient navigation, even in novel circumstances^1^. This concept gained momentum with the discovery of ‘place cells’ in the hippocampus, neurons that fire selectively at specific positions in the environments^2–4^. Since then, the neural underpinnings of cognitive maps have been studied extensively, revealing a vast body of knowledge about the firing properties of neurons that comprise cognitive maps in the brains of rodents^5^, primates (including humans)^6–17^, and other animals^18,19^.

These foundational studies indicate that the hippocampus not only captures features of the environment, but also the relationships between them and the animal’s actions within it. For example, many hippocampal neurons carry information in the form of activity that is greatest at a particular location in the environment (the cell’s ‘place field’)^2^, while others store information not only about place, but also about contextual features such as the animal’s movement direction^20,21^, running speed^20^, or movement history^22,23^. Hippocampal neurons can also learn to represent more abstract spaces, such as the position in a sound landscape^24^, accumulated evidence^25^, arbitrary relationships between concepts, objects, or events^11,26,27^, and other non-spatial dimensions^24,28,29,25,30–32^. Despite an extensive body of knowledge about the neuronal firing properties constituting hippocampal cognitive maps, and recent ideas concerning their algorithmic structure^33–37^, we are still yet to fully characterize the formation of cognitive maps during the entire learning phase of moderately complex tasks. Acquiring such empirical data is crucial, as it facilitates identification of the learning principles that drive the creation of cognitive maps. While cognitive maps have been shown to form rapidly in novel environments and modify rapidly in response to changes in familiar environments^38–40^, technical limitations such as the limited number of total recorded neurons (hundreds) and short durations (days) of longitudinal tracking of individual neurons have impeded the ability to study the formation of cognitive maps representing more complex relationships that require extensive exploration and learning. Here, we leveraged technological advances that allowed us to follow neural activity in thousands of hippocampal neurons in each mouse, stably for many days or weeks, as they learned to perform a task requiring them to form cognitive maps representing spatial, temporal, and abstract relationships as they interacted with a complex but predictable environment.

Our results show that during learning, mice proceed through a stereotypical series of behavioral changes that are mirrored by structured changes in neural activity. Specifically, hippocampal activity undergoes a series of decorrelation steps that orthogonalize neural activity in regions of the environment where sensory stimuli are similar, but task demands differ. We analyzed the representational structure of the task, visualized the low-dimensional geometry of the neural activity, and compared neural activity in the hippocampus to unit activity in a variety of cognitive models and artificial neural networks trained using the same task structure. We show that day-to-day dynamics of hippocampal activity are consistent with the formation of a state machine, consisting of orthogonalized latent representations of task-specific features. The transitions between these latent states, each encoding specific task features or segments, are learned to predict the dynamics of the animal’s interaction with the environment. Remarkably, orthogonal states can represent similar sensory stimuli over learning, highlighting latent task structure as the driver of state orthogonalization.

We show further that this orthogonalized state machine (OSM) can be reproduced by a type of Hidden-Markov-Model (HMM) model called a Clone-Structured Causal Graph (CSCG)^35,41^ or by a biologically plausible recurrent neural network (RNN) trained using Hebbian learning^42^, an unsupervised local learning rule that does not require backpropagation of error^43^. Despite their effectiveness in sequence modeling, we show that gradient-based learning of models such as LSTMs^44^ or Transformers^45^ do not naturally generate representational geometries that mirror those observed during animal learning. We further demonstrate that neural activity shows flexible usage of the OSM in altered tasks conditions such as the introduction of new visual cues and adjustment in the lengths of track segments. In sum, these findings shed light on the architecture and learning principles governing the formation of cognitive maps and provide potential guides for the design of future artificial systems.

## RESULTS

### Learning the 2-alternative cue-delay-choice task

We trained transgenic mice expressing GCaMP6f in hippocampal neurons to navigate in a virtual reality (VR) environment while head-fixed to enable imaging of neural activity as they learned the relationship between visual cues and the future location of water reward delivery in two linear tracks (Fig. 1a; see Methods for details). On each trial, water was delivered at one of two reward zones, either near or far from the beginning of the track, which we call R1 and R2, respectively. Prior to these rewarded locations, a visually distinct indicator cue (Ind) perfectly predicted the rewarded location (Fig. 1a, bottom). Efficient execution of this 2-alternative cue-delay-choice task (2ACDC) requires mice to form and use long-term memory of the relationship between the indicator cues and reward locations and short-term memory of the indicator cue after it disappears and before the rewarded location.

**Figure 1.**
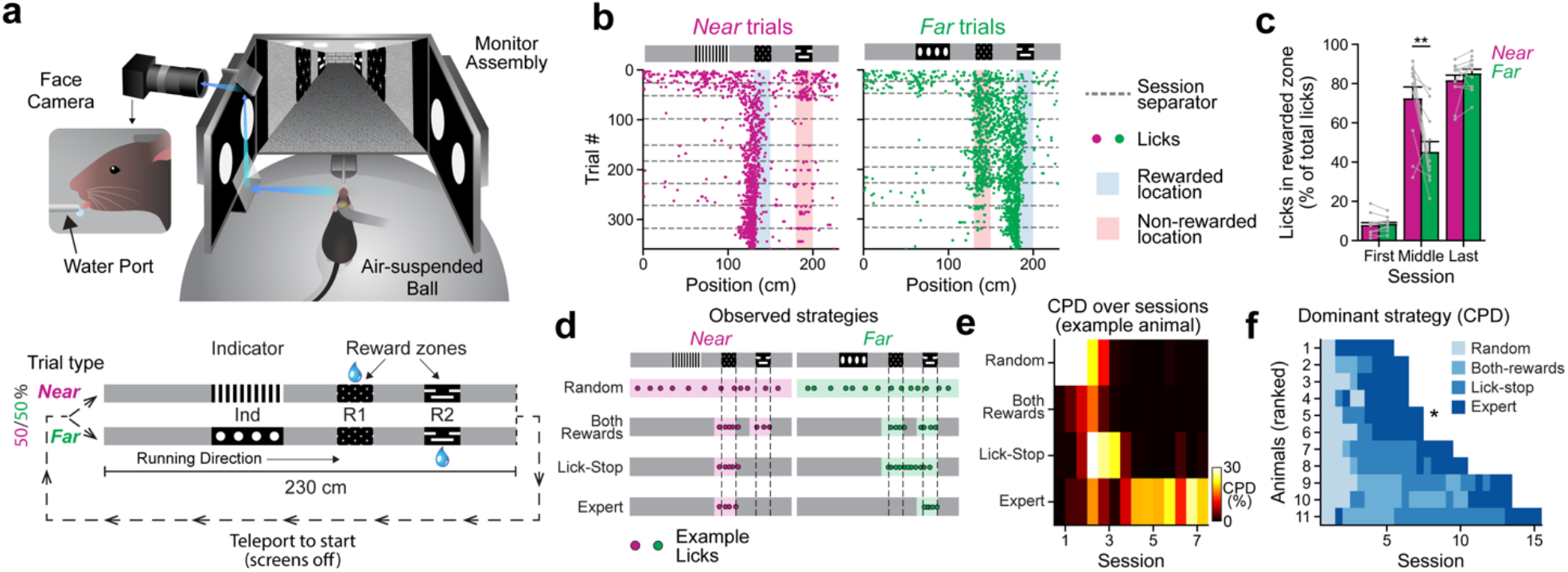
Animals exhibit stage-wise behaviors when learning the 2-alternative cue-delay-choice task. **(a)** Diagram of the virtual reality behavior setup (top) and illustration of the task (bottom). Ind: Indicator regions, R1: Near reward zones, R2: Far reward zones. **b)** Example lick patterns of a single animal across sessions for both the *Near* and *Far* trial types. Each dot represents a single lick within a trial and dashed lines indicate separation between daily sessions. **(c)** Percentage of licks at the correct reward locations for both the *Near* and *Far* trial types for the first session, an intermediate session, and the last session in which the animals show expert performance (n = 11 mice; *P* <0.01**, dependent samples t-test). **(d)** Illustration of four different behavior strategies animals exhibit during learning. Colored shadings denote the non-zero regions within the basis functions for regression against licking density across track locations. **(e)** Coefficients of Partial Determination for the four behavior strategies for one example animal. **(f)** Dominant strategy for all animals over sessions (ranked by learning speed). Asterisk denotes the animal shown in e. Values in e and f are calculated by splitting each session into two parts of equal duration (start/end). Number of sessions per strategy, Random: 1.9 ± 0.7, Both-rewards: 2.0 ± 2.5, Lick-stop: 3.3 ± 2.8, Expert: 3.6 ± 0.5.

Mice were initially trained for 5 days (one-hour session each day) to run on a spherical treadmill and collect randomly delivered water rewards in the dark. Subsequently, screens were turned on to display the VR environment. In each subsequent one-hour daily session, mice performed ∼80-200 trials (124 ± 43 trials, n = 11 mice), with the reward location depending on the trial type. Both *Near* and *Far* trial types were presented sequentially in a randomized manner (Methods). To initiate a new trial, mice had to run to the end of the corridor, which was decorated by a ‘brick wall’ cue, and a 2-second period of dark screens (‘teleportation’) preceded the next trial. The two trial types shared identical visual cues at all locations other than the indicator region. Outside of the indicator region and the reward regions, the walls of the virtual corridor were decorated with relatively featureless gray wood grain (‘gray’ regions). This sensory ambiguity within trials (four gray regions) and across trials (both gray regions and reward-zone cues are visually identical) is a key feature of the task. For the first 1-3 days of training, water rewards were delivered even if the mouse did not lick in the rewarded zone, until consistent anticipatory licking was observed. On all subsequent days, mice were rewarded with a drop of water on any trial only if they licked in the correct reward zone. No penalty was imposed for licking in other locations. Thus, mice learned the task through exploration, presumably motivated to slow down and lick only when rewards were expected.

We assessed learning by plotting licking behavior as a function of position for all trials across several days of training on the 2ACDC task (Fig. 1b). Initially, mice licked throughout the entire track, but they quickly learned to restrict licking to portions of the track near the two reward zones in both trial types. This change in behavior occurred within 2-3 sessions in all animals (Suppl. Fig. 1) Around the same time, mice developed an intermediate strategy and learned to suppress licking after receiving a reward. As a result, licking behavior was near optimal for the *Near* trial type (not licking at the far reward zone), but remained suboptimal for the *Far* trial type (Fig. 1c, middle session), because licking began at the near reward zone and was often sustained until the reward delivery at the far reward zone. With additional training, mice eventually learned to suppress licking in the near reward zone on the *Far* trial type, thus achieving close to optimal performance on both trial types (Fig. 1c, last session). Thus, licking behavior appears to evolve in stages, marked with distinct behavioral strategies (Fig. 1d): (1) random licking, (2) licking in both reward locations, (3) licking in reward locations and stop licking until a reward is collected (‘lick-stop’), and (4) only licking near the correct reward locations (‘expert’). We used a statistical method called Coefficient of Partial Determination (CPD) to assess the contribution of each of the four behavior strategies to the overall behavior of the animals. Using these four behavior strategies as regressors accounted for 36.5 ± 5.9 % of variance in licking behavior averaged across all sessions (n = 9 ± 3 sessions per mouse). This explained variance percentage is within the expected range for behavioral studies involving complex tasks^46^. By removing a regressor corresponding to each behavior strategy one at a time, we were able to determine its unique contribution to the behavior. CPD analysis demonstrated that these four strategies emerged in successive waves at different points during the learning process (Fig. 1e), which was mirrored by gradual changes in the profile of animals’ running speed (Suppl. Fig. 2). Despite variations in the number of sessions required for different animals to reach expert performance, the progression through these dominant behavior strategies was remarkably consistent (Fig. 1f).

### Longitudinal imaging of hippocampal activity during learning

Prior to training, all mice were implanted with a cranial window to enable imaging of neural activity using GCaMP6f expressed in pyramidal neurons in area CA1 of the dorsal hippocampus (Fig. 2a; see Methods). Activity was imaged using a two-photon random access mesoscope (2P-RAM)^47^. The 3-mm cranial window was readily imaged with the 2P-RAM, which has a 5-mm field of view (FOV). Thousands of cells (within single sessions: 4682 ± 827 on average per animal across all sessions, range: 3813 - 6490; maximum single session cell count: 5545 ± 848 on average per animal, range 4266 - 7309. n = 11 mice) mostly near the center of the FOV, were readily resolved and reidentified in each session across several weeks of training and imaging (cells tracked across sessions: 3954 ± 661 on average per animal, range 3034 - 5354; see Methods. Fig. 2b; Suppl. Fig. 3, 4).

**Figure 2.**
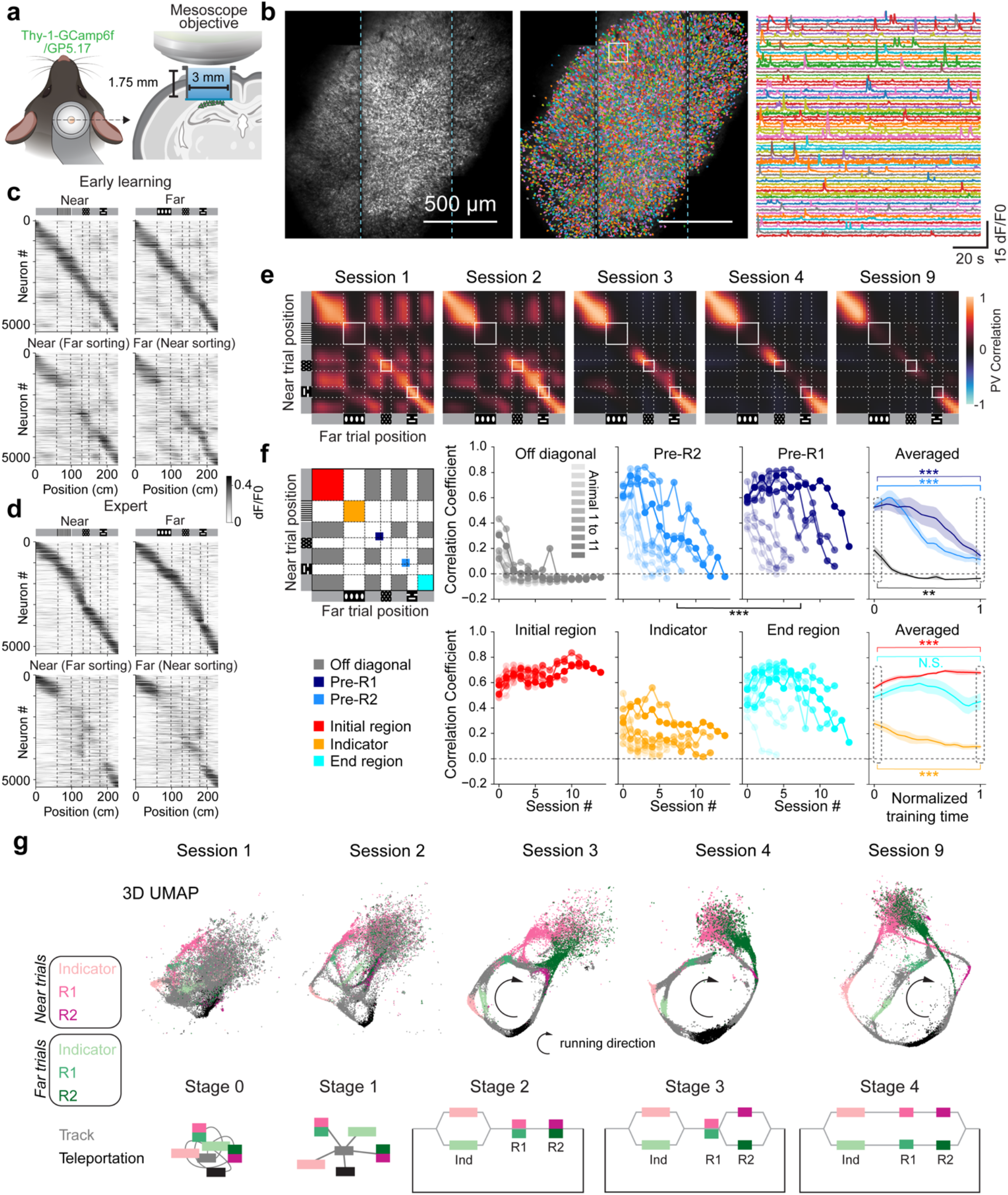
Progressive decorrelation of neural activity during learning. **(a)** Diagram of the window implant for CA1 imaging. **(b)** Example field of view, cell segmentations, and example extracted fluorescent signals. Cyan dashed lines mark the boundary between vertical scanning stripes of the 2P-RAM microscope. **(c)** Trial-averaged neural activity versus track position for both the *Near* and *Far* trial types on session 2 in a representative animal (Animal 7 in Fig. 1f). Top row: cells ordered by the center of mass (COM) of the largest place field by indicated trial type. Bottom row: cells ordered by opposite trial type. **(d)** Similar to **c**, but for session 9 of the same animal. **(e)** *Near*-versus-*Far* PV Cross-correlation matrices along all track positions for sessions 1, 2, 3, 4, 9 for the same example animal. **(f)** PV correlation averaged for different regions on the cross-correlation matrix across sessions, for the off-diagonal gray region correlations (gray), Pre-R2 region (light blue), and Pre-R1 region (dark blue), Initial region (red), Indicator region (orange), End region (cyan) shown for each animal separately and averaged across all of them. Comparing all sessions, a significant difference was observed between the Pre-R1 and Pre-R2 regions (Wilcoxon signed-rank test, *P* < 0.001***). Comparisons between the first and last session revealed significant changes in the Pre-R2, Pre-R1, Indicator (decreasing over sessions), and Initial region (increasing over sessions) with *P* < 0.001***. For the off-diagonal region, a significant decrease in correlation was also observed with *P* < 0.01**. In contrast, changes in the End region were not significant (N.S.). **(g)** 3D UMAPs and corresponding state diagrams of the neural manifold for 5 different sessions across learning from the same example animal in e, the 2D views of the UMAP are shown here were chosen to best illustrate the learning dynamics captured by the 3D structure.

By the second session, many cells exhibited elevated activity at well resolved positions along the track. Ordering cells according to the position of the peak activity revealed a clearly resolvable diagonal band of spatial responses tiling the whole virtual track in both trial types (Fig. 2c). Ordering cells according to their spatial activity pattern for the opposite trial type also showed a diagonal spatial band indicating many cells are active at similar locations in both trial types. However, activity differed between trial types most prominently at the indicator cue position (Fig. 2c), indicating that sensory information dominates the neural activity at this early stage. Indeed, cells that were most active in one of the four gray regions also showed moderate activity in other gray regions (Fig. 2c). After several more days of training, these neural activity-based ‘maps’ of the 2ACDC task increasingly differentiated within single trial types (among multiple gray regions) and between the two trial types (Fig. 2d).

### Stage-wise changes of hippocampal activity during learning

The representational structure of the *Near* and *Far* trial types was compared by computing a population vector (PV) correlation for the two trial types (Methods), which decreased systematically in selected positions during training. Analysis of the cross-correlation between *Near* and *Far* trial types across all regions of the track indicated that the indicator cue region had the lowest correlation, as expected from the differences in visual stimuli at this location (Fig. 2e, f). Both within and between trial types, the four gray regions of the track were moderately correlated on the first session for all animals but by the third session, the correlation had significantly reduced (Fig. 2e, f & Suppl. Fig. 5 showing PV angles also approaching 90°), suggesting that the hippocampus had orthogonalized its representations of these visually similar regions. Between-trial-type correlations decreased in an ordered, stage-like fashion, with the neural activity at the track region right before the far rewarded zone (Pre-R2) decorrelating generally earlier than the region before the near rewarded zone (Pre-R1; Fig. 2e, f). Neural activity corresponding to the indicator cue, while low from the onset, decorrelated further with increased exposure to the two trial types (Fig. 2e, f). While most track regions underwent complete decorrelation into near orthogonal representations, the beginning and end of the track remained correlated throughout training for most animals (Suppl. Fig. 6). This suggests that the decorrelation process is shaped by the task structure, as after collecting the reward, the animal has no information on which trial type it will be in next until it sees the next indicator.

Taken together, the initial decorrelation of gray regions suggests that the hippocampus first learns the sequential structure of the two linear environments by learning to differentiate each of the four visually similar gray regions within the linear tracks. During training, the hippocampus learns to further differentiate the similar visual cues between the two trial types, following a highly consistent sequential order, as indicated by the progressive decorrelation towards orthogonalization of neural activity across specific track locations: Initially, based presumably on whether the animal successfully collected reward at the near reward zone (rewarded in *Near*, but not yet rewarded in *Far*), neural activity decorrelates immediately after the animal has passed the first reward region. As training progresses, the remaining correlated activity prior to the first reward region eventually undergoes orthogonalization. This gradual decorrelation in neural activity co-evolves with the animal’s progressively improving licking behavior (Suppl. Fig. 8).

### The hippocampal representation has properties of an orthogonalized state machine

We further visualized the day-to-day dynamics of neural activity utilizing a non-linear dimensionality reduction technique, specifically Uniform Manifold Approximation and Projection (UMAP)^48^. A single embedding space was used to reduce the activities of thousands of cells to points in a low-dimensional (3D) UMAP space, using longitudinally registered data gathered across all days of imaging, where each point represents the activity of all cells in a single imaging frame (see Methods, Fig. 2g). Notably, this UMAP representation not only echoed the gradual decorrelation and orthogonalization we previously described (Fig. 2e, f and Suppl. Fig. 5, 6), but it also allowed us to intuitively observe the overall topological changes of the neural manifold during learning.

Here we describe the UMAP from a representative animal, exhibiting all learning stages. The UMAP representation from the initial session distinctively clusters the neural activity associated with each sensory cue. Despite this differentiation, the overall neural manifold appears relatively unstructured at this stage (Fig. 2g, Stage 0). By the second day, the UMAP adopts a ‘hub-and-spoke’ appearance (Fig. 2g, Stage 1), with the hub corresponding to all gray regions and the spokes corresponding to activity trajectories between a gray region and all other cues (i.e., indicator cue, near and far reward cues, and the dark teleportation region). This structure hints that the neural activity associated with the concept of ‘linear tracks’ may not be fully developed at this point. An additional scattered point cloud near the reward-zone embeddings corresponds to periods of water-reward licks and a post-reward period where the mouse was not running. By the third session, the UMAP adopts a ring-like structure that is closed by activity during the 2-second dark teleportation period linking the end of one trial and the start of the next trial (Fig. 2g, Stage 2). As training progresses, the activity trajectories for the two trial types become increasingly distinct, eventually resembling a split-shank wedding ring, consisting of a band that splits into two strands with a diamond in the center. Here, the splitting band corresponds to the principal manifold of neural activity while the mouse is running and the diamond corresponds to the point cloud when the animal is stationary, mostly during and right after reward consumption. We speculate that this reward-associated point cloud, observable in the UMAP representations at all stages (Suppl. Fig. 7), may be related to replay of neural activity and may contribute to synaptic plasticity in the hippocampus as well as its downstream targets^49^. The gradual appearance of a split-shank ring UMAP mirrors the dynamics of trial-type decorrelation described above (Fig. 2e, f).

The observed progressive changes in the representational structure reflected by both the correlation matrices and UMAPs resemble a gradually evolving state machine undergoing several meaningful intermediate stages and finally reaching a structure capturing the essence of the task (Fig. 1a, bottom and Fig. 2g, state diagrams below the UMAPs). This process involves several stages of disambiguation of similar sensory inputs at different regions along the two trial types in the populational activity level, eventually producing orthogonalized state representations for previously latent states of the task. We call this learned manifold an “orthogonalized state machine” (OSM).

The orthogonalization process of neural representations, as described earlier, reflects changes in the firing properties of individual neurons that occur together with the changes in an animal’s behavior. As training progresses, the firing properties of numerous neurons undergo modifications. One prominent change in the early stages of learning involves neurons tuned for multiple gray regions becoming more selective, firing at fewer gray regions and in many cases a single one (Fig. 3a). As learning continued, neurons displayed increasingly distinct tuning across the *Near* and *Far* trial types in regions including the pre-reward track regions (“pre-R1” and “pre-R2”). This encompassed neurons that were initially silent but later became active in specific regions for one trial type and not the other, as well as neurons that started as active on both trial types but eventually became trial-type-specific by decreasing the activity in the other trial type (Fig. 3b, c). We found that such ‘splitter cells’^22,23,50^ were generated at every stage of learning (Fig. 2e, Fig. 3). In summary, these single-cell tuning changes described above can be understood from the perspective of the hippocampus progressively extracting the latent task structure from ambiguous sensory experiences.

**Figure 3.**
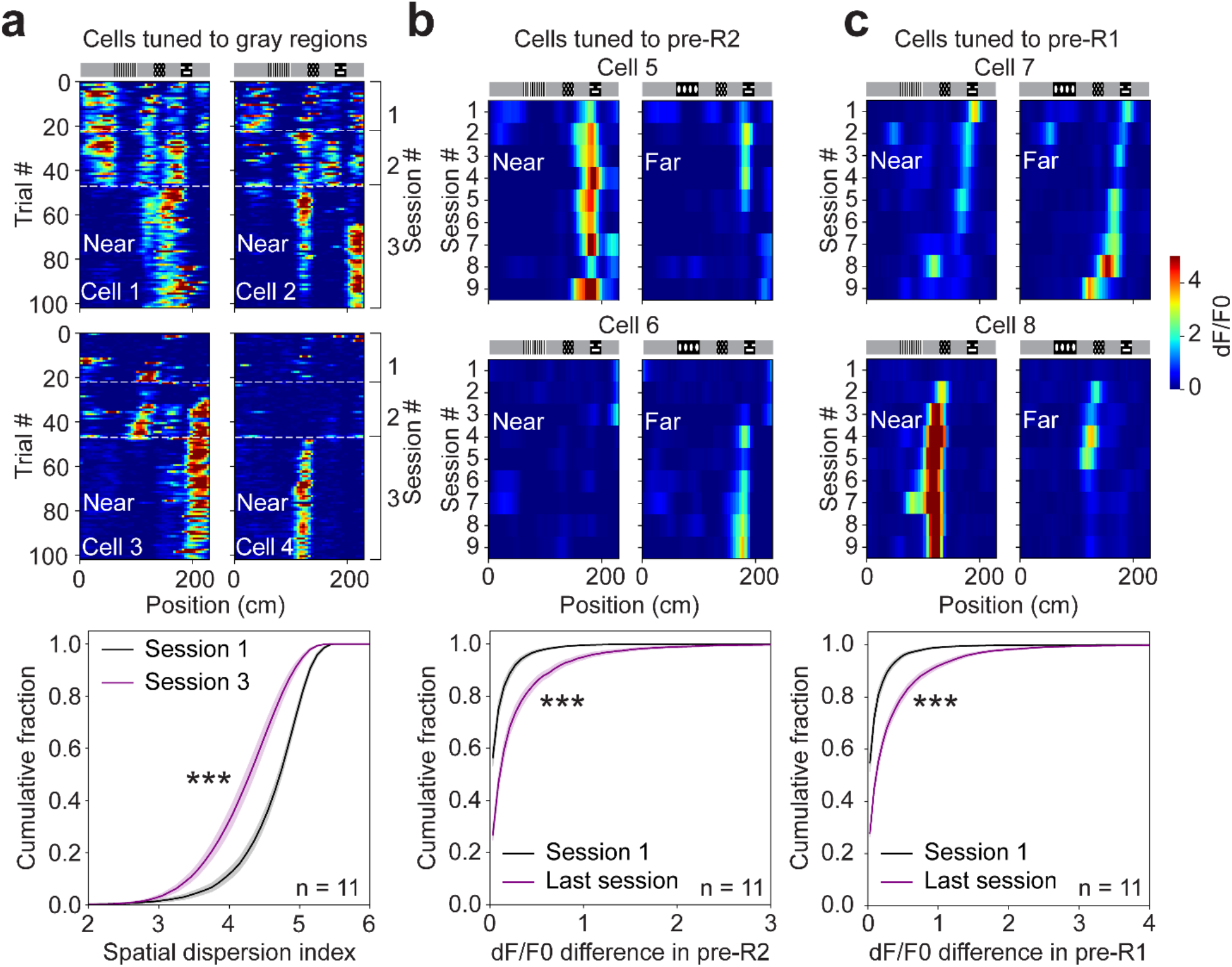
Single-cell tuning changes during learning. **(a)** Top: positional tuning for 4 example cells tuned to gray regions from one animal across *Near* trials for stage 1 to 2 transition (first 3 sessions for this example animal). Bottom: cumulative percentage plot of the spatial dispersion index (entropy of normalized spatial tuning, see Methods) for all cells with gray region tuning within the first 3 sessions in the *Near* trial for session 1 and session 3 (Wilcoxon rank-sum test, *P* < 0.001***). **(b)** Top: session-averaged positional activity for the *Near* and *Far* trials across 9 sessions for two example cells tuned to pre-R2 region for either *Near* or *Far* trials. Bottom: cumulative percentage of the dF/F0 difference in pre-R2 between *Near* and *Far* trial types for all cells with significant tuning in pre-R2, for session 1 and last session (session 9) (Wilcoxon rank-sum test, *P* < 0.001***). **(c)** Similar to **b**, but for the pre-R1 region (Wilcoxon rank-sum test, *P* < 0.001***).

### Comparing the structure of hippocampal maps to mathematical and computational models

The large number of neurons we recorded over many days of training presents a unique opportunity to probe the learning algorithms that can lead to the gradual emergence and the final representational structure of cognitive map of the 2ACDC task. Aligned with previous work^34,35,37,41,51–57^, we conceptualize the problem as learning the underlying structure of the world by learning to predict sequences of sensory inputs.

To explore the class of models that can recapitulate animals’ learning process, we first used an HMM-based model called the ‘clone-structured causal graph’ (CSCG)^35^. Fundamentally, HMMs and their variants like CSCGs aim to uncover hidden structures from sequential data, capturing meaningful latent states and their temporal dependencies. CSCGs consist of ‘clones’ that correspond to states with occupancy probabilities that are influenced by current and past sensory stimuli that are experienced sequentially. The model is constructed using the Expectation Maximization (EM) algorithm^58^ to predict the sequence of sensory symbols representing different track regions and reward delivery (as a sensory cue) in the 2ACDC task (Fig. 4b, Methods, Suppl. Fig. 9, 10; Methods). The resulting clones and their occupancy probabilities can be compared to neural activity in the hippocampus by conceptualizing them as units and activity. Consistent with previous work^35^, we found that units in the CSCG trained using sequences from the 2ACDC task progressed through stages of orthogonalization that closely mirrored what we observed in the hippocampus (Fig. 4b, Suppl. Fig. 9). For example, the CSCG gradually learns to use different clones to represent visual cues with similar sensory features arriving in different contexts (e.g., gray cue following the indicator cue or the first reward cue). In addition, the CSCG generated using the learned transition matrix progressed through various stages during its training that bear a striking resemblance to the representations of neural activity we observed in mice (Fig. 2e, g; Suppl. Fig. 9d). The finding that CSCG, a cloned HMM, can recapitulate significant aspects of animal learning indicates that extracting meaningful latent states from potentially ambiguous, ongoing sequential sensory experiences is a key feature of hippocampal learning.

**Figure 4.**
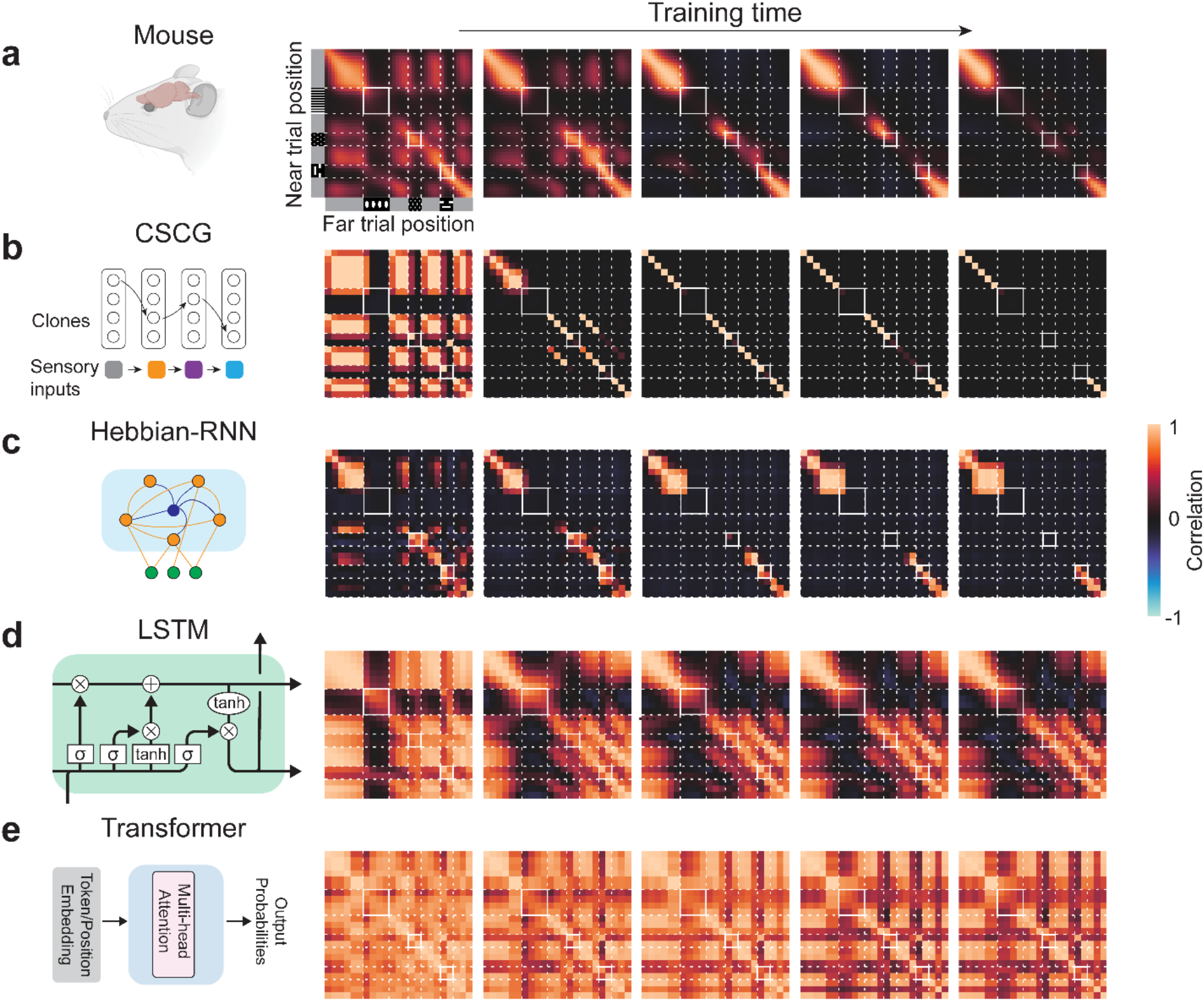
Representational structure during learning for mice and different models. **(a)** CA1 PV cross-correlation between *Near* and *Far* trials in an example mouse over 5 different sessions (Sessions 1, 2, 3, 4, 9, from left to right, same data from Fig. 2e). **(b-e)** Cross-correlation matrices for different models trained to learn from sensory sequences sampled from a structurally similar task to 2ACDC: **(b)** CSCG, (c) RNN with Hebbian plasticity (Hebbian-RNN), (d) LSTM, (e) Transformer. CSCG and Hebbian-RNN are trained until convergence; LSTM and Transformer start with random performance and trained until expert performance (Suppl. Fig. 12).

Prior work has suggested that Hebbian plasticity such as spike-timing-dependent plasticity (STDP)^59,60^ can result in attractors that stably encode sequences^61,62^ in a manner that is robust to noise^63,64^, form predictive maps^65–67^, and approximate HMM learning^51,68^. To test this idea further, we built a spiking RNN model that included a soft winner-take-all (WTA) mechanism, which leverages the principle of feedback inhibition to ensure that only the highest firing neurons remain active within the network. Using only a timing-based Hebbian plasticity rule^51^ based on local activity (i.e., no end-to-end training), the model (Hebbian-RNN) progressed through stages of decorrelation to result in an orthogonalized representation of the 2ACDC task that is qualitatively similar to the CSCG trained using expectation maximization (Fig. 4c). However, in Hebbian-RNN, pre-R1 decorrelated before pre-R2, whereas in CSCG, and in the hippocampus, pre-R2 decorrelates before pre-R1. These results suggest that Hebbian plasticity and feedback inhibition in a RNN are sufficient to reproduce various aspects of single cell tuning changes (Suppl. Fig. 11) and a final representation of the 2ACDC task with features like the OSM representation we observed in the hippocampus, but that additional mechanisms will be required to fully mimic the learning dynamics (see Discussion).

Artificial neural networks such as LSTMs^44^ and Transformers^45^ are powerful models for sequence learning. Using gradient descent via backpropagation of error, we trained an LSTM model to predict the next input in a sensory sequence (reward delivery also as a sensory cue) in a 2ACDC task (Fig. 4d; see Methods). Although the LSTM model could learn to perform perfectly on the task (Suppl. Fig. 12), unit activity in the well-trained model differed strikingly from the properties of neural activity we observed in the mouse hippocampus. Specifically, most of the activity corresponding to the same sensory input in different latent states remain highly correlated (Fig. 4d) and trial-type differences in activity in the well-trained agent were minor compared to the activity levels (Suppl. Fig. 12). Using a similar approach, we used gradient descent to train a decoder-only Transformer to learn to predict the next input in the 2ACDC task. Again, while the trained model accurately predicted future states in the 2ACDC task, unit activity in the model differed markedly from what we observed in mice, with properties like those of the LSTM model (Suppl. Fig. 12).

Next, we examined if adding various regularization schemes could help LSTMs to produce representational structures that more closely resemble those observed in the brain. Our findings suggested that commonly employed regularization techniques — those that promote sparseness or introduce noise during the training phase — were unsuccessful in capturing the orthogonal representational structure in the fully trained model (Suppl. Fig. 12). Notably, however, we could coerce the model into generating an orthogonalized representational structure by using a cost function that specifically penalized non-zero values in the correlation matrix between hidden states for both *Near* and *Far* trials. This is consistent with the recognized capability of approximating complicated functions in neural networks^69,70^ (Suppl. Fig. 12f).

### Novel task features lead to adaptation of the existing hippocampal state machine

To investigate how the hippocampal state machine, once learned by a mouse, can be utilized when new elements are introduced, we expanded and later modified the structure of the task. First, after mice learned the task with the original indicator cues (cue pair A) we replaced them with two unfamiliar visual patterns. To do this we developed four unique indicator pairs (cue pair B, C, D, and E) and presented them to mice that had already learned the original cue pair (Fig. 5a). Every day, the mice were initially exposed to cue pair A for a duration of 5 to 10 minutes, after which the indicators for the task were replaced with one of the novel pairs. This change enabled us to collect neural activity data for both the original and new cue pairs. Training on the new cue pair continued until the mouse could proficiently execute the task, demonstrated by restricting its licking to the rewarded location and just before it on 75% of the trials for three successive sessions. Mice were then sequentially trained on each subsequent novel indicator pair on the following days in the same manner. Through this training process mice learned the new cue sets in significantly fewer trials (147 ± 39 trials for the new cue sets compared to 483 ± 70 trials for the original cue set; n = 3 mice; *P* < 0.05*, unpaired *t*-test. Fig. 5b).

**Figure 5.**
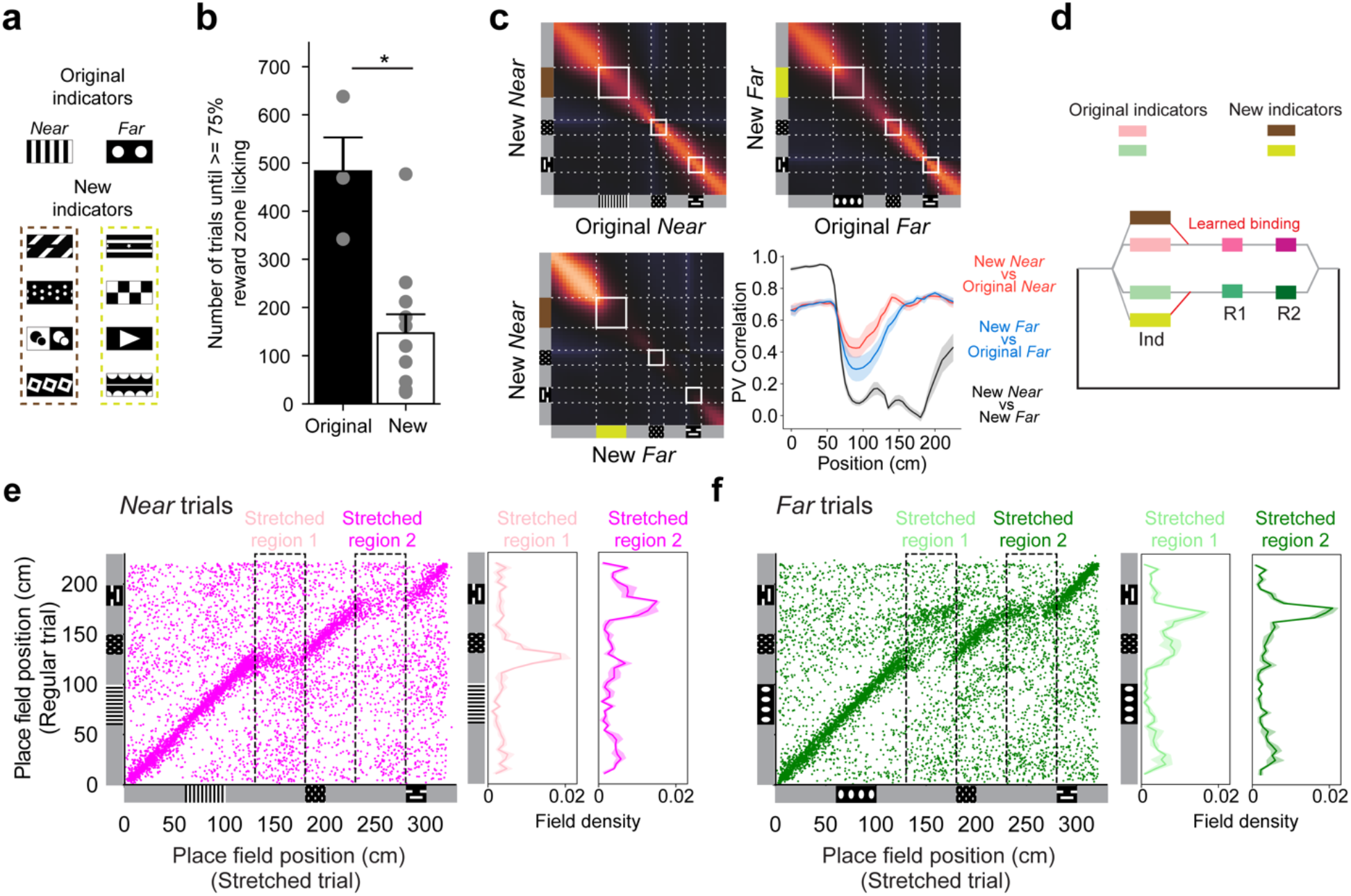
State machines can be flexibly used in novel settings. **(a)** Original and novel indicator pairs. **(b)** Number of trials before high performance (restricting licking to the rewarded location and just before it on 75% of the trials) for each indicator pair (bar graph showing mean ± s.e.m, p = 0.039, unpaired *t*-test). Dots represent data from a single animal for one indicator pair. **(c)** PV cross-correlation between the neural activity for trials with the original and new indicators for the same trial type (*Near*, top left; *Far*, top right); between *Near* and *Far* trials for the new indicators (bottom left). Bottom right: quantification of the diagonal cross-correlations in 3 cross-correlation matrices. **(d)** Conceptual diagram for the incorporation of new indicator states into an existing state machine. **(e)** Place field locations in the stretched *Near* trials plotted against those in the regular *Near* trials (n = 3 mice, data pooled together). The histograms of field locations in the two stretched regions are plotted to the right. **(f)** Similar to **e** but for the *Far* trials.

Comparing PV correlations of the neural activity for trials with novel indicators versus the original indicators revealed high similarities in neural representations between the old and new tasks in all track regions except the indicator region (Fig. 5c). In other words, the neural activity in the presence of new indicators mirrored the common task structure while maintaining information about the visual identities of the novel indicator cues. In terms of the state machine framework, this suggests that once a state machine is established in the hippocampus, it can be effectively utilized and reused for new task variants. New task elements can be integrated into the existing state machine, either through the creation of new states or linking new sensory inputs to existing states (Fig. 5d). This flexible adaptation and integration, in turn, expedites learning.

In a second variation of the task, we extended the length of the gray zones following the indicator cue and after the first reward cue, thus requiring animals to travel longer distances to reach the reward zones (Fig. 5e, f; Methods). These ‘stretched trials’ were introduced every 5-6 trials, to evaluate how well-trained animals for the original task respond to the altered environment without extensively training them to adjust to these task modifications (Methods). In both stretched *Near* (*Near’*) and stretched *Far* (*Far’*) trial types, mice displayed a tendency to lick towards the beginning of the usual reward location, even though the reward cue was not yet encountered (Suppl. Fig. 13a). A comparison of the tuning location for each cell during normal and stretched trials of the same type (i.e., *Near*-*Near’* or *Far*-*Far’*) provided insights into how the animals might perceive the modified task (Fig. 5e, f). As expected, for the initial part of the tracks (ranging from the start of the track to the gray region just prior to the stretched region), which are identical to the original trials, the place fields were tuned to similar locations between the regular and stretched trials (Fig. 5d). As the animal enters the first stretched region in the *Near* trials, which is the first deviation from the normal trial, many cells tuned for the gray region before the first reward location (pre-R1) were active throughout the stretched region (Fig. 5e, n = 3 mice). This potentially indicates that the animal may believe it remains in the same patch of gray zone right before the first reward region. Interestingly, during *Far* trials, the difference in reward location expectation drives a different pattern to emerge. When the animal enters the first stretched region in far trials, the place cells didn’t persistently extend their activity. Instead, they rapidly shifted their tuning to align with the region just before the second reward area (pre-R2), as if the animal is anticipating the second reward location (Fig. 5f). However, when the animal eventually sees the first reward cue, the representation quickly resets and anchors to the representation reflecting the first reward region in the original task. These results imply that, when the animals encounter modified components of the task, the neural representations can settle into discrete states to mirror inferred latent states under conditions of uncertainty (Fig. 5e, f). In conclusion, these discoveries substantiate the idea that the learned cognitive map exhibits the properties of a state machine that can potentially infer and flexibly use learned states in novel situations.

## DISCUSSION

We assessed neural activity in a large population of neurons as mice learned a complex task over the course of several days to weeks. The results indicate that a cognitive map of the task appears gradually in the hippocampus in parallel with improvements in task performance. Well-trained mice exhibit robust short-term and long-term memory—processes that are consistent with the structure of the mature cognitive map, including the ability to use the map to produce effective behavior in novel environments with similar structure and altered features. The cognitive map has features of a state machine with orthogonalized representations of latent states that the animal must discover to perform the task efficiently. Computational modeling suggests that many features of this orthogonalized state machine (OSM), including the gradual dynamics of its formation, share properties with a type of Hidden Markov model (HMM) called a clone-structured causal graph (CSCG)^35^ that is learned using the expectation maximization (EM) algorithm. We show further that a recurrent neural network (RNN) coupled with winner-take-all (WTA) dynamics and trained using Hebbian plasticity can mimic many features of the OSM and its formation. This combination of RNN, WTA, and Hebbian plasticity has been previously shown to mimic EM^51^, highlighting that known biological mechanisms can construct graph-like representations of environments where animals repeatedly experience sequences of sensory stimuli, including rewards that are delivered in latent contexts. Strikingly, we find that widely used sequence learning models in AI, trained to predict the next element in a sequence by using backpropagation of error, do not naturally produce orthogonalized representations like those found in the mouse hippocampus.

These results have important implications for understanding the plasticity mechanisms that may contribute to the formation of the hippocampal cognitive maps. Foremost among them, our modeling indicates that Hebbian plasticity in a RNN with WTA dynamics is sufficient to construct the map. This form of plasticity does not require feedback from other brain areas, as it is determined entirely by the relative timing of pre and postsynaptic spikes local to the modified synapse. This is fundamentally different from the methods typically used to adjust synaptic weights in artificial neural networks, which rely on explicitly defined cost functions^71^. Nevertheless, there is good evidence that feedback-based plasticity is important in the hippocampus^72,73^ and it has been proposed to be a key element in approximating the backpropagation of error algorithm in the brain^74^. Thus, we do not claim that Hebbian plasticity operates on its own, but rather, that Hebbian plasticity is likely to be one important mechanism for the formation of cognitive maps. Feedback-based mechanisms involving target, error, or reward signals are also likely to be instrumental. These mechanisms may work in tandem with Hebbian plasticity to construct cognitive maps and/or they may be more involved in refining behavior policy and other task-specific functions by selectively routing information from the established cognitive maps to other brain regions mediating behavioral policies. Our data indicate that task representations and behavioral policies based upon them are formed in lockstep, as suggested by previous theory^75^. A likely candidate mechanism for the contribution of synaptic plasticity during feedback is behavioral time scale synaptic plasticity (BTSP)^76^.

Despite the well-known existence of synaptic plasticity in CA1, plasticity mechanisms in other brain regions are likely to underpin our observations as well. For example, CA1 gets most of its excitatory synaptic input from CA3, where recurrent connections^77^, attractor dynamics^78^, and Hebbian plasticity have all been observed^79^. Plasticity in CA3, as well as other brain regions, may thus result in changes in the firing of the pyramidal neurons we imaged in CA1.We propose that the existence of multiple forms of synaptic plasticity across different brain regions allows unsupervised and supervised (and reinforcement) learning to work together to reduce sensory interference, build robust models of the environment, and direct the content of these models to promote adaptive behaviors. Identifying the loci and molecular mechanisms of these processes is a central challenge for neuroscience.

A classical concept in computer science, a finite state machine is a computational structure consisting of a finite set of states with the transitions between them based on defined inputs or conditions^80^. States reflect current sensory input from the environment and the animal’s own body, as well as latent information such as the recent history of sequential observations. Transitions are constrained by the current state and neurally encoded transition probabilities, and determined by the animal’s own movements and the sensory input it receives from the environment. We posit that neural activity in the hippocampal OSM could shape behavior, such as speeding up, slowing down, or licking. These behaviors in turn influence neural activity, and thus transitions to new states in the hippocampal OSM, both by changing the animal’s external and internal sensory experience and by changing the stimuli coming from the environment. The hippocampal OSM operates in closed loop with the rest of the brain, the animal’s body, and its environment to produce the properties of a state machine.

Our findings are related to and expand upon many previous studies, including those describing the concepts of partial remapping^81^ (upon modifications of the environment, only some neurons change their spatial firing patterns while other neurons remain unchanged), rate remapping^82^ (the location of place cell tuning remains unchanged but the firing rate varies), and global remapping^82–84^ (the activity patterns in two different environments are not correlated), which refer to subtle and more extensive changes in the hippocampal code when the animal’s environment changes subtly or dramatically^85^. Our study indicates that learning the 2ACDC task can gradually induce partial remapping, with cells tuned to specific track segments (e.g., reward cues) showing progressively decorrelated neural activity, while other cell populations, specifically those tuned to the beginning or end of the track, sustain highly correlated activity across trial types. Our results also suggest that rate remapping and partial remapping could be intermediate stages for the gradual orthogonalization process of two contexts^86,87^, as the PV correlation for various parts of the track decreases gradually from highly correlated to near orthogonal representations as learning progresses. Additionally, we observed that the visually distinct indicators for both trial types show highly decorrelated neural activity from the very beginning of task exposure, which further deepens and approaches orthogonalization as learning progresses. This suggests that even representations of distinct sensory cues could be further orthogonalized if they are associated with different task states.

Our work is also related to previous reports of “splitter cells” as well as the concept of pattern separation, as both constitute forms of the decorrelated representations. Splitter cells (and related phenomena) are neurons that exhibit differential firing when similar sensory cues are experience in different contexts^22,23,29,88,89^. Our study extends this idea from differential neuronal tuning of specific locations to a systematic orthogonalization process that generates cells matching the core features of splitter cells at each stage of the learning process (e.g., gray regions, and two separate reward zones). The emergence of such generalized splitter cells suggests that an important learning mechanism and computation performed by the hippocampus are intrinsically linked to the separation of similar sensory experiences based on their different task implications^22,29,34,35,40,89^. These observations are related to pattern separation, which is the ability of animals to distinguish between subtly different sensory cues by representing small differences^90–94^. Our data suggests that different sensory cues already show high levels of decorrelation early in learning, which might be the result of receiving already pattern-separated input from upstream regions (e.g., DG). Such differences in the sensory features of the indicator cues can be further decorrelated when the task benefits from it. While pattern separation can be enabled by expansion recoding^95^ or neuronal nonlinearities in recurrent networks (without weight changes)^96^, our experimental data and modeling suggest that local Hebbian plasticity likely plays an important role. Such computations allow the hippocampus to approximate the environment’s generative process through the creation of hippocampal ‘state cells’ representing the gradually discovered task-related latent states during each learning stage^35,37,97–100^.

Sparse orthogonal representations have long been proposed by others to constitute a powerful mechanism for memory and intelligence^36,101–110^. However, we find that traditional AI systems trained with backpropagation of error—specifically LSTMs and Transformers—do not naturally produce the sparse orthogonalized representations. However, we found that this key property of the hippocampal OSM could be observed in LSTM when the cost function explicitly penalized activity correlation between the two trial types. Influential work in AI has shown that the enforced-orthogonalization approach is a powerful tool in self-supervised learning systems^111^. In addition, the ability to tailor LSTMs to exhibit orthogonalized states has implications for uncovering mechanisms that implement error backpropagation in the brain^74^. Other frameworks for modeling hippocampal function include predictive coding and representations^112–114,33,37,53^, statistical learning^108,115^, latent states inference^34,35,98,116–118^, compression^36,52,119,120^, and hippocampal models tasked with generalization such as the Tolman Eichenbaum Machine^97^ and spatial memory pipeline^121^. These varied approaches reflect the rich complexity of the hippocampal system, and each offers distinctive insights into our understanding of cognitive maps and the broader mechanisms of learning and memory in the brain.

Several avenues for future research emerge from our findings. While both CSCG and the online Hebbian-RNN can capture the orthogonalized representational structure, they differ in how they mirror the process by which this orthogonalization unfolds in animals. Specifically, the Hebbian-RNN model accurately reproduces the end-stage representation but deviates in the exact sequence of orthogonalization when compared to CSCG and observed neural dynamics of the animal. This discrepancy highlights an opportunity for future research to develop more refined and biologically realistic models of hippocampal cognitive maps, for example by incorporating plasticity mediated by forward and backward replay mechanisms^122,123^. Moreover, a compelling direction would be to juxtapose the cognitive maps of the hippocampus with those of the neocortex. Given their roles as complementary learning systems^124^, the hippocampus and neocortex are thought to capture distinct facets of the external world. While the hippocampus has been posited to store memories of specific, idiosyncratic experiences, the neocortex leans toward storing more generalizable memories^124–126^. Hence, co-recording the cognitive maps from both brain regions during learning sessions could provide invaluable insights into their interactions and respective contributions to learning and behavior.

In summary, in this study we provide detailed, comprehensive, longitudinal data on the formation of a cognitive map in the hippocampus during learning of a moderately complex task, which progresses in stages over the course of several training sessions. The excellent match between CSCGs, Hebbian-RNNs, and hippocampal data suggest that state machines with sparse, orthogonalized representations are likely to provide a powerful framework for neural computation^104^, memory^105,127^, and intelligence^128^. These results also underscore the need to understand how different learning rules such as local Hebbian learning and gradient descent-based learning might synergistically function to promote a more effective and efficient learning process in both natural and artificial learning systems^71,129,130^.

## METHODS

All procedures were performed in accordance with the Janelia Research Campus Institutional Animal Care and Use Committee guidelines. Both male and female GCaMP6f (Thy1-GCaMP6f^131^) transgenic mice were used, 3 to 6 months old at the time of surgery (3 to 8 months old at the beginning of imaging studies).

### Surgery

Mice were anesthetized with 1.5 - 2.0% isoflurane. A craniotomy on the right hemisphere was performed, centered at 1.8 mm AP and 2.0 mm ML from the bregma using a 3 mm diameter trephine drill bit. The overlying cortex of the dorsal hippocampus was then gently aspirated with a 25-gauge blunt-tip needle under cold saline. A 3 mm glass coverslip previously attached to a stainless-steel cannula using optical glue was implanted over the dorsal CA1 region. The upper part of the cannula and a custom titanium headbar were finally secured to the skull with dental cement. Animals were allowed to recover for a minimum of 2 days before being put under water-restriction (1.0-1.5ml/day), in a reversed dark-light cycle room (12-hour light-dark cycle).

### Behavior

#### VR setup

The VR behavior setup was based on a design previously described in Cohen et al. 2017^132^. The cylindrical treadmill consisted of a hollowed-out Styrofoam ball (diameter = 16 inches, 65g) air-suspended on a bed of 10 air-cushioned ping-pong balls in an acrylic frame. Animals were head-fixed on top of the treadmill using a motorized holder (Zaber T-RSW60A; Optics Focus MOG-130-10 and MOZ-200-25) with their eyes approximately 20 mm above the surface. To translate the movement of the treadmill into VR, two cameras separated at 90 degrees were focused on 4 mm^2^ regions of the equator of the ball under infrared light^132^. Three axis movement of the ball was captured by comparing the movement between consecutive frames at 4kHz and readout at 200 Hz^132^. A stainless-steel tube (Inner diameter= 0.046 inches), attached to a three-axis motorized stage assembly (Zaber NA11B30-T4A-MC03, TSB28E14, LSA25A-T4A, and X-MCB2-KX15B), was positioned in front of the animal’s mouth for delivery of water rewards. The animal was shown a perspective corrected view of the VR environment through 3 screens (LG LP097QX1 with Adafruit Qualia bare driver board) placed roughly 13 cm away from the animal (Fig. 1a). This screen assembly could be swiveled into position using a fixed support beam. All rendering, task logic and logging was handled by a custom software package called Gimbl (https://github.com/winnubstj/Gimbl) for the Unity game engine (https://unity.com/). All inter-device communication was handled by a MQTT messaging broker hosted on the VR computer. Synchronization of the VR state with the calcium imaging was achieved by overlaying the frame trigger signal of the microscope with timing information from inbuild Unity frame event functions. To observe the mouse during the task without blocking its field-of-view, we integrated a periscope design into the monitor assembly. Crucially, this included a 45° hot-mirror mounted at the base of a side monitor that passed through visible light but reflected IR light (Edmund optics #62-630). A camera (Flea3-FL3-U3-13Y3M) aimed at a secondary mirror on top of the monitor assembly could hereby image a clear side view of the face of the animal. Using this camera, a custom Bonsai script^133^ monitored the area around the tip of the lickport and detected licks of the animal in real time that were used in the VR task as described below.

#### Head fixation training

After recovering from surgery, mice were placed on water restriction (1.0 -1.5 ml per day) for at least 2 weeks before starting any training. Body weight and overall health indicators were checked every day to ensure animals remained healthy during the training process. Mice were acclimated to experimenter handling for 3 days by hand delivering water using a syringe. For the next 3 sessions, with the VR screens turned off, mice were head-fixed on the spherical treadmill while water was randomly dispensed from the lick-port (10 ± 3 sec interval; 5 µl per reward). These sessions lasted until animals acquired their daily allotment of water or until 1 hour had passed. We observed that during this period most mice started to run on their own volition. Next, we linked water rewards to bouts of persistent running and increased this duration across sessions till the animal would run for at least 2 seconds straight (∼5 sessions). During this time, we also slowly increased the height of the animal with respect to the treadmill surface across sessions to improve performance. Animals that did not show sufficient running behavior to acquire their daily allotment of water were discarded in further experiments.

#### 2-alternative cue-delay-choice (2ACDC) task

At the beginning of each trial mice were placed at the start of a virtual 230 cm corridor. The appearance of the walls was uniform except at the location of three visual cues that represented the indicator cue (40 cm long) and the two reward cues (near/far; 20 cm long). Depending on the trial type, a water reward (5 µl) could be obtained at either the near, or far reward cue (near and far reward trials). The only visual signifier for the current trial type was the identity of the indicator cue at the start of the corridor. For the first 2-3 sessions the animal only had to run past the correct reward cue in order to trigger reward delivery (‘Guided’ sessions). On all subsequent sessions mice had to lick at the correct reward cue (‘Operant’ sessions). No penalty was given for licking at the incorrect reward cue. In other words, if the animal licked at the ‘near’ reward cue during a ‘far’ trial type then the animal could still receive a reward at the later ‘far’ reward cue. Upon reaching the end of the corridor the VR screen would slowly dim to black, and mice would be teleported to the start of the corridor to begin the next semi-randomly chosen trial with a 2-second duration. The probability of each trial type was 50% but in order to prevent bias formation caused by very long stretches of the same trial type, sets of near/far trials were interleaved with their number of repeats set by a random limited Poisson sampling (lambda= 0.7, max repeats=3). The identity of the indicator cue was kept hidden for the first 20 cm of the trial and was rendered when the animals pass the 20 cm position. To internally track the learning progress of the animal we utilized a binarized accuracy score for each trial depending on whether the animal only licked at the correct reward cue. Once the animal had 3 sessions where the average accuracy was above 75%, we considered the animal to have learned that cue set.

#### 2ACDC task with novel indicators

For 3 mice out of the 11 well-trained animals on the original 2ACDC task, we subsequently trained them to perform the 2ACDC task novel indicator cues. After reaching 3 consecutive sessions with >75% task accuracy for the original 2ACDC task, the novel task was introduced the following session, but with the original task shown for the first 5 to 10 minutes at the beginning of each session before switching completely to the new task. When the animal can perform the new task for 3 consecutive days > 75%, we move on to the next novel indicator pair until the last one is finished (4 novel indicator pairs in total).

#### 2ACDC task with extended gray regions

As another modification to the original task design, the gray regions were extended in certain trials which we call the “stretched trials”. In the stretched trials, the linear track was extended from 230 cm to 330 cm, and the reward positions were moved from [130, 150] cm to [180, 200] cm (the 1st rewarding (Near) object), and [180, 200] cm to [280, 300] cm (the 2nd rewarding (Far) object). Note that the distance between the indicator cue and the Near object in the stretch trial is equal to the one between the indicator cues and the Far object in the normal trial. During a session with stretch trials, following a 5-minute warm-up using only the normal 2ACDC trial, the stretch trial was adopted at intervals of every 5 or 6 trials.

### Calcium Imaging

Neural activity was recorded using a custom-made 2p-RAM mesoscope^47^ and data acquired through ScanImage software^134^. GCaMP6f was excited at 920 nm (Chameleon Ultra II, Coherent). 3 adjacent ROIs (each 650 µm wide) were used to image dorsal CA1 neurons. The size of the ROIs was adjusted to ensure a scanning frequency at 10 Hz. Calcium imaging data were saved into tiff files and were processed using the Suite2p toolbox (https://www.suite2p.org/). This included motion correction, cell ROIs, neuropil correction, and spike deconvolution as described elsewhere^135^.

### Multiday alignment

To image the same cells across subsequent days, we utilized a combination of mechanical, optical, and computational alignment steps (Suppl. Fig. 3, 4). First, mice were head fixed using a motorized head bar holder (see above) allowing precise control along three axes (roll, pitch, and height) with submicron precision. Coordinates were carefully chosen at the start of training to allow for unimpeded movement and reused across subsequent sessions. The 2p-RAM microscope was mounted on a motorized gantry, allowing for an additional three axis of alignment (anterior/posterior, medial/lateral, and roll). Next, we utilized an optical alignment procedure consisting of a ‘guide’ led light that was projected through the imaging path, reflected off the cannula cover glass and picked up by a separate CCD camera (Suppl. Fig. 3b). Using fine movement of both the microscope and the head bar, the location of the resulting intensity spot on the camera sensor could be used to ensure exact parallel alignment of the imaging plane with respect to the cover glass.

To correct for smaller shifts in the brain tissue across multiple sessions, we took a high-resolution reference z-stack at the start of the first imaging session (25 μm, 1 μm interval; Suppl. Fig. 3c). The imaging plane on each subsequent session was then compared to this reference stack by calculating a cross correlation in the frequency domain for each imaging stripe along all depth positions. By adjusting the scanning parameters on the remote focusing unit of the 2p-RAM microscope we finely adjusted the tip/tilt angles of the imaging plane to achieve optimal alignment with the reference stack. We used a custom online Z-correction module (developed by Marius Pachitariu^136^, now in ScanImage), to correct for Z and XY drift online during the recording within each session, using a newly acquired z-stack for that specific session.

To find cells that could be consistently imaged across sessions we first performed a post hoc, non-rigid, image registration step using an averaged image of each imaging session (Diffeomorphic demon registration; python image registration toolkit) to remove smaller local deformations (Suppl. Fig. 3g-i). Next, we performed hierarchical clustering of detected cells across all sessions (jaccard distance; Suppl. Fig. 4). Only putative cells that were detected in 50% of the imaging sessions were included for further consideration. We then generated a template consensus mask for each cell based on pixels that were associated with this cell on at least 50% of the sessions. These template masks were then backwards transformed to the spatial reference space of each imaging session to extract fluorescence traces using Suite2p.

## Data analysis

### Coefficient of Partial Determination

To assess the unique contribution of each behavioral strategy (random licking, licking in both reward locations, lick-stop, and expert) to overall animal behavior, we used the coefficient of partial determination (CPD). In this analysis, a multivariable linear regression model was first fitted using all behavioral strategies as regressors, providing the sum of squares error (SSE) of the full model (SSE_fullmodel_). Each regressor was then sequentially removed, the model refitted, and the SSE without that regressor (SSE_∼i_) was computed. The CPD for each regressor, denoted as CPD_i_, was then calculated as CPD_i_ = (SSE_∼i_ - SSE_fullmodel_) / SSE_∼i_, revealing the unique contribution of each behavioral strategy to the overall variance in licking behavior.

### Place field detection

To identify significant place cells, we utilized an approach based on Dombeck et al., 2010^137^ (but see also Grijseels et al., 2021^138^ for overall caveats with such approaches). Place fields were determined during active trials, indicated by active licking within reward zones, and at running speeds greater than 5cm/s. For detecting activity changes related to position we first calculated the calcium signal by subtracting the fluorescence of each cell mask with the activity in the surrounding neuropil using suite2p. Next, the baseline fluorescence activity for each cell was calculated by first applying gaussian filter (5 secs) followed by calculating the rolling max of the rolling min (‘maximin’ filter, see suite2p documentation). This baseline fluorescence activity (F0) was used to calculate the differential fluorescence (dF/F0), defined as the difference between fluorescent and baseline activity divided by F0. Next, we identified significant calcium transient event in each trace as events that started when fluorescence deviated 5σ from baseline and ended when it returned to within 1σ of baseline. Here, baseline σ was calculated by binning the fluorescent trace in short periods of 5 seconds and considering only frames with fluorescence in the lower 25^th^ percentile.

Initially, putative place fields were identified by spatially binning the resulting ΔF/F0 activity (bin size: 5 cm) as continuous regions where all dF/F0 values exceeded 25% of the difference between the trial’s peak and the baseline 25^th^ percentile dF/F0 values. We imposed additional criteria: the field width should be between 15-120 cm in virtual reality, the average ΔF/F0 inside the field should be at least four times greater than outside; and significant calcium transients should occur at least 20% of the time when the mouse was active within the field (see above). To verify that these putative place fields were not caused by spurious activity we calculated a shuffled bootstrap distribution for each cell. Here, we shuffled blocks of 10 secs calcium activity with respect to the animal’s position and repeated the same analysis procedure described above. By repeating this process 1000 times per cell we considered a cell to have a significant place field if putative place fields were detected in less than 5% of the shuffles.

### Population vector (PV) analysis

For the analysis of similarity of representation between *Near* vs *Far* trial types, we performed population vector correlation on the fluorescence dF/F0 data. Each 5cm spatial bin, we define the population vector as the mean dF/F0 value for each neuron. Fluorescence data were included only when the animal’s speed exceeded 5cm/sec. The cross-correlation matrix was generated by calculating the Pearson correlation coefficient between all location pairs across the two trial types.

### Spatial dispersion index

To evaluate the extent of spatial dispersion in place tuning across single cells – such as distinguishing between tuning to single positions versus multiple positions, we take the single cell tuning curve over track positions and normalize it so the area under the curve is 1. The spatial dispersion index is defined as the entropy of this normalized dF/F0 signal by: Entropy = - ∑ [p(i) * log2 p(i)], where p(i) denotes the probability associated with each position bin index.

### Uniform Manifold Approximation and Projection (UMAP)

To visually interpret the dynamics of high-dimensional neural activity during learning, we utilized UMAP on our deconvolved calcium imaging data. The UMAP model was parameterized with 100 nearest neighbors, 3 components for a three-dimensional representation, and a minimum distance of 0.1. The ’correlation’ metric was used for distance calculation. The data, a multidimensional array representing the activity of thousands of cells concatenated from several imaging sessions, was fitted into a single UMAP model. This resulted in a three-dimensional embedding, where each point characterizes the activity of the neuron ensemble at a single imaging frame.

## Modeling

### CSCG

In the 2ACDC task, the combination of position along the track and trial type defines a state of the world (*z*). Although this state is not directly observable to the animal, it influences the sensory observation (*x*) that the animal perceives (Suppl. Fig. 9a). The sequence of states in the environment obeys the Markovian property, whereby the probability distribution of the next state (i.e., next position and trial type) depends only on the current state, and not all the previous states, assuming the animal always travels at a fixed speed. When an animal learns the structure of the environment and builds a map, it tries to learn which states (position, trial type) are followed by which states, and what sensory experience they generate. This can be viewed as a Markov learning problem. A Hidden Markov model (HMM) consists of a transition matrix whose elements constitute *p*(*z*_*n*−1_ ∣ *z*_*n*_) i.e., the probability of going from state *z*_*n*_ at time *n* to *z*_*n*−1_ at time *n* + 1, an emission matrix whose elements constitute *p*(*x*_*n*_∣ *z*_*n*_), ie. the probability of observing *x*_*n*_when the hidden state is *z*_*n*_, and the initial probabilities of being in a particular hidden state *p*(*z*_1_).

CSCG (Cloned Structured Causal Graph) is an HMM with a structured emission matrix where multiple hidden states, referred to as clones, deterministically map to the same observation. In other words, *p*(*x*_*n*_ = *j*|*z*_*n*_ = *i*) = 0 if *i* ∉ *C*(*j*) and *p*(*x*_*n*_ = *j*|*z*_*n*_ = *i*) = 1 if i∈ *C*(*j*), where *C*(*j*) refers to the clones of observation *j* ^35^ (Suppl. Fig. 9b). The emission matrix is fixed and CSCG learns the task structure by only modifying the transition probabilities (Suppl. Fig. 9b), making the learning process more efficient. Baum Welch expectation maximization algorithm was used to update the transition probabilities such that it maximizes the probability of observing a given sequence of sensory observations^139–141^.

We trained CSCG on sequences of discrete sensory symbols mimicking the sequence of patterns shown to the animals in the 2 tracks. Each 10 cm segment of the track was represented by a single sensory symbol. Additionally, the teleportation region was represented by a distinct symbol repeated thrice, spanning 30cm. In the rewarded region, the animals could receive both visual input and a water reward simultaneously. However, our model could only process a single discrete stimulus at a time. Thus, we divided the rewarded region into two parts. We presented the visual cue first, mimicking the animals’ ability to see the rewarded region ahead before reaching it. Subsequently, we presented a symbol representing the water stimulus, which was shared across the two trials. The near trial sequence, denoted as [1,1,1,1,1,1,2,2,2,2,1,1,1,4,6,1,1,1,5,5,1,1,7,0,0,0], and the far trial sequence, denoted as [1,1,1,1,1,1,3,3,3,3,1,1,1,4,4,1,1,1,5,6,1,1,7,0,0,0]’, were used. Where 1 represented the gray regions, 2 and 3 the indicators for near and far tracks respectively, 4 the visual observation associated with the first reward zone, 5 denoted the visual stimulus associated with the far reward zone, 6 represented the common water reward received in both tracks, 7 the brick wall at the end of each trial, and 0 indicated the teleportation region (Suppl. Fig. 9c).

We initialized the model with 100 clones for each sensory observation symbol and performed 20 iterations of the expectation-maximization process at each training step with sequences from 20 randomly selected trials, comprising both near and far trial types. We extracted the transition matrix at different stages of learning and used Viterbi training algorithm to refine the solution^35^ and then plotted the transition matrix as a graph, showing only the clones that were used in the representation of the 2 trials (Suppl. Fig. 9d). We ran multiple simulations and compared how correlation between the two trial types changed over learning for different positions along the track (Suppl. Fig. 9e).

We also explored alternate sequences of sensory stimuli. In one variant, we provided the water symbol prior to the visual symbol of the reward zone (for ex. […1,1,1,6,4,1,1,1…] where 6 represents the water and 4 the visual symbol). Additionally, we introduced a symbol that conjunctively encoded the simultaneous water reward and visual symbol (for ex. and […111,4,6,111…] in the near trial and […111,5,8,111…] in the far trial, where 6 denotes a combined code for water and visual R1, and 8 represents a combined code for water and visual R2) (Suppl. Fig. 10a). While the final learned transition graphs matched for all the 4 sequence variants, the exact sequence of learning differed. Specifically, reward cue followed by a visual cue for reward zone often led to decorrelation of Pre-R1 followed by Pre-R2 (Suppl. Fig. 10b, c), contrary to what is often observed during learning in animals. Therefore, for further analysis, we utilized the visual followed by common reward sequence, as it led to a learning trajectory that closely matched the learning observed in animals.

### Hebbian-RNN

Work by Kappel et al. (2014)^51^ showed that a local Hebbian learning rule in a recurrent neural network (RNN) can approximate an online version of HMM learning. We used an RNN consisting of *K* = 100 recurrently connected neurons and *N* = 96 feedforward input neurons. The feedforward input neurons carried orthogonal inputs for each of the 8 sensory stimuli, with 12 different neurons firing for each stimulus (Suppl. Fig. 11a). The recurrent weights *V* and feedforward weights *W* were initialized from a normal distribution with 0 mean and standard deviation 2.5 and 3.5 respectively. The membrane potential of the *k*^*th*^ neuron at time *t* is given by 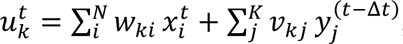, where *w*_*ki*_ is the feedforward weight from input neuron *i* to RNN neuron *k*, *v_kj_* is the recurrent weight from neuron *j* to neuron *k*, Δ*t* = 1*ms* is the update time, and 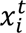 and 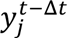 are exponentially filtered spike trains of the feedforward and recurrent neurons respectively (exponential kernel time constant, 20 ms). The probability of neuron *k* firing in Δ*t* was computed by exponentiating the membrane potential and normalizing it through a global inhibition, 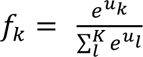 For each neuron *k*, spikes were generated with a probability of *f_k_* by a Poisson process, with a refractory period of 10 ms during which the neuron cannot spike again. When the post-synaptic neuron *k* spiked, then the weights onto neuron *k* were updated as 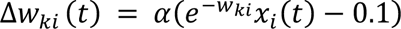 and 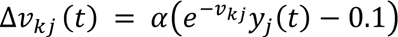, where α is the learning rate (0.1) and *y_j_*(*t*) is exponentially filtered spike train. Both weights *V* and *W* were kept excitatory. We computed the correlation between the RNN representation of different positions in the near and far trial types at different stages during learning and compared it with cross-correlation matrices for animals.

### LSTM

We utilized LSTMs for predicting the next sensory symbol in sensory sequences in a 2ACDC task. Task sequences incorporated numerical symbols with unique meanings: ’1’ denoted the gray region; ’2’ and ’3’ represented near and far cues respectively; ’4’ and ’5’ indicated near and far reward cues; ’6’ symbolized reward; and ’0’ denoted teleportation. An example of a near trial followed the structure: 1,1,1,1,1,1,2,2,2,2,1,1,1,4,6,1,1,1,5,5,1,1,0, and far trial follows the structure: 1,1,1,1,1,1,3,3,3,3,1,1,1,4,4,1,1,1,5,6,1,1,0. We converted these numerical symbols into one-hot encodings to represent these categories. The LSTM model was constructed with PyTorch, having 128 neurons in the single hidden layer and eight neurons in the output layer. For optimization, we used Adam with a learning rate of 3e-4 with a Cross Entropy Loss. Initially, the model was trained without any regularization to establish a baseline performance. Later, we implemented different regularization techniques, namely L1, L2, dropout^142^, and correlation penalization (akin to Barlow Twins^111^), in distinct simulation versions (Suppl. Fig. 12).

### Transformers

We implemented a Transformer architecture based on the minGPT repository (https://github.com/karpathy/minGPT). We designed trials that represent sequences in an environment with each symbol denoting a specific meaning similar to the LSTM simulations. We created sequences of trials with random starts for a total of 3000 batches. The trials were randomly assembled in a sequence length of 10 trials. We then generated sequences of ’near’ and ’far’ trials and selected random 100-element chunks to form our input sequences. The vocabulary size and block size were set according to our dataset. The Transformer model was trained using a learning rate of 3e-4 for a total of 600 iterations. The objective was to predict the next sensory symbol, using the Cross Entropy Loss, similar to the LSTM. During test time, 4-symbol sequences were used to test the trained model’s next-input prediction accuracy.

## ACKNOWLEDGMENTS

We thank Hessameddin Akhlaghpour, Antonio Fernandez-Ruiz, Brad Hulse, Albert Lee, Brett Mensh, Gabriela Michel, Anja Payne, Sandro Romani, Yuhan Wang, and Lin Zhong for their comments on the manuscript. We thank Marius Pachitariu for the assistance with mesoscope imaging pipelines. We thank Albert Lee and Jae Sung Lee for their technical guidance on CA1 window surgeries. We thank Vasily Goncharov and Dmitri Tsyboulski for mesoscope technical support. We thank Gabriela Michel, Boaz Mohar, Yuhan Wang, Xinyu Zhao, and other current and former members of the Spruston lab for their discussion, technical assistance, and feedback throughout the project. We thank Salvatore Dilisio and Sarah Lindo for their assistance in animal surgeries and the Janelia Vivarium team for animal support. We thank Matt Botvinick, Zeb Kurth-Nelson, Dharsh Kumaran, Kimberly Stachenfeld, and Jane Wang from DeepMind for discussions regarding AI models. We thank François Chollet, Luke Coddington, Ian Cone, Josh Dudman, Stephano Fusi, Mehrdad Jazayeri, Jim Knierim, Sam Lewallen, Jeffrey Magee, Brett Mensh, Andrew Saxe, and Yaniv Ziv for valuable discussions. We thank the Janelia Experimental Technology team, including Jon Arnold, Bruce Bowers, Tobias Goulet, Dan Smith, Steve Sawtelle, and Alex Sohn for technical assistance. We thank Julia Kuhl for the illustration of the VR setup. This work was supported by the Howard Hughes Medical Institute.

## SUPPLEMENTARY FIGURES

**Supplementary figure 1.**
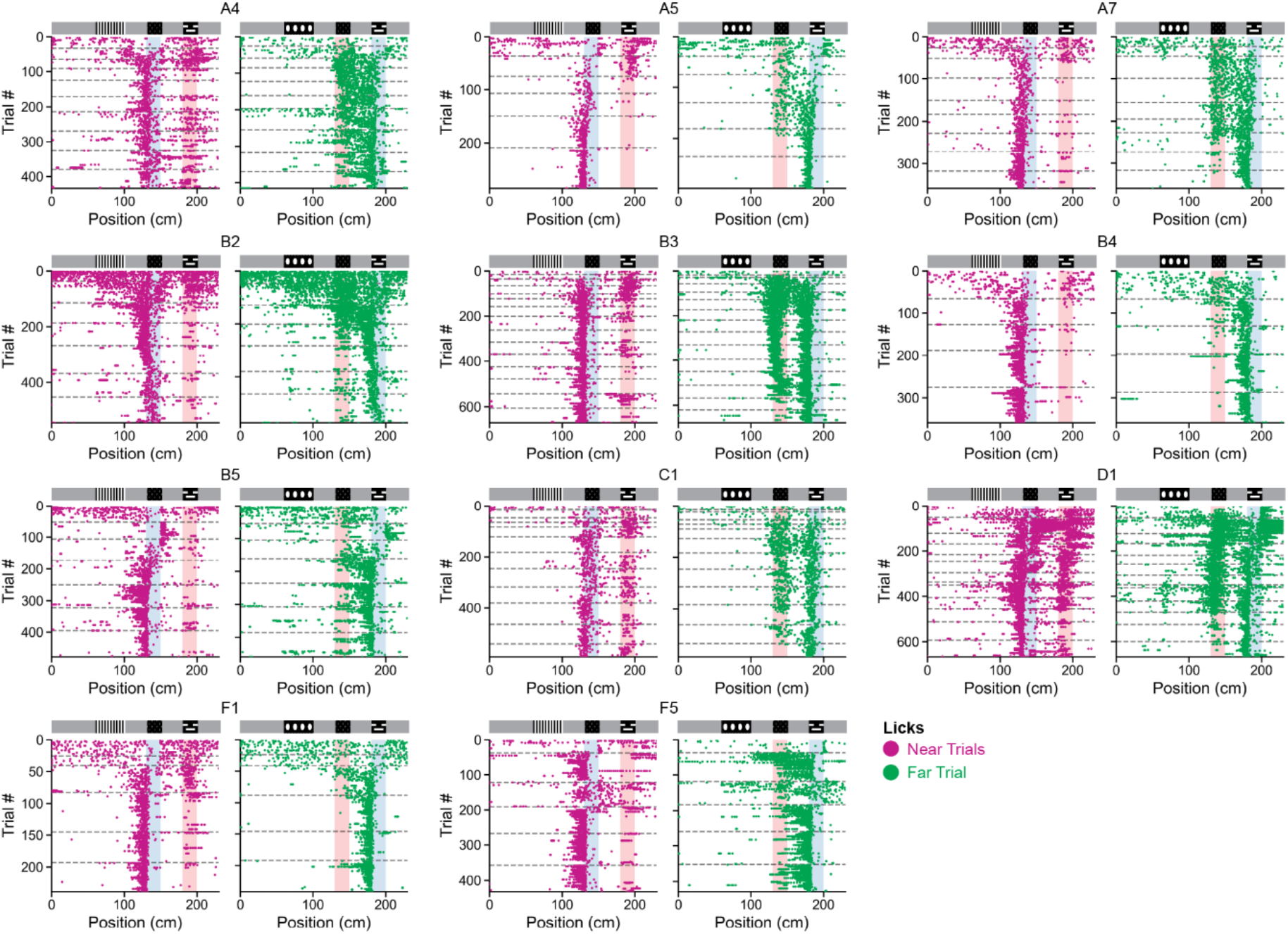
Changes in licking behavior during learning across all animals. Each dot indicates a single lick made by an animal relative to its position on the track in either a ‘near’ (magenta) or ‘far’ trial type (green). Dashed lines indicate the end of a session.

**Supplementary figure 2.**
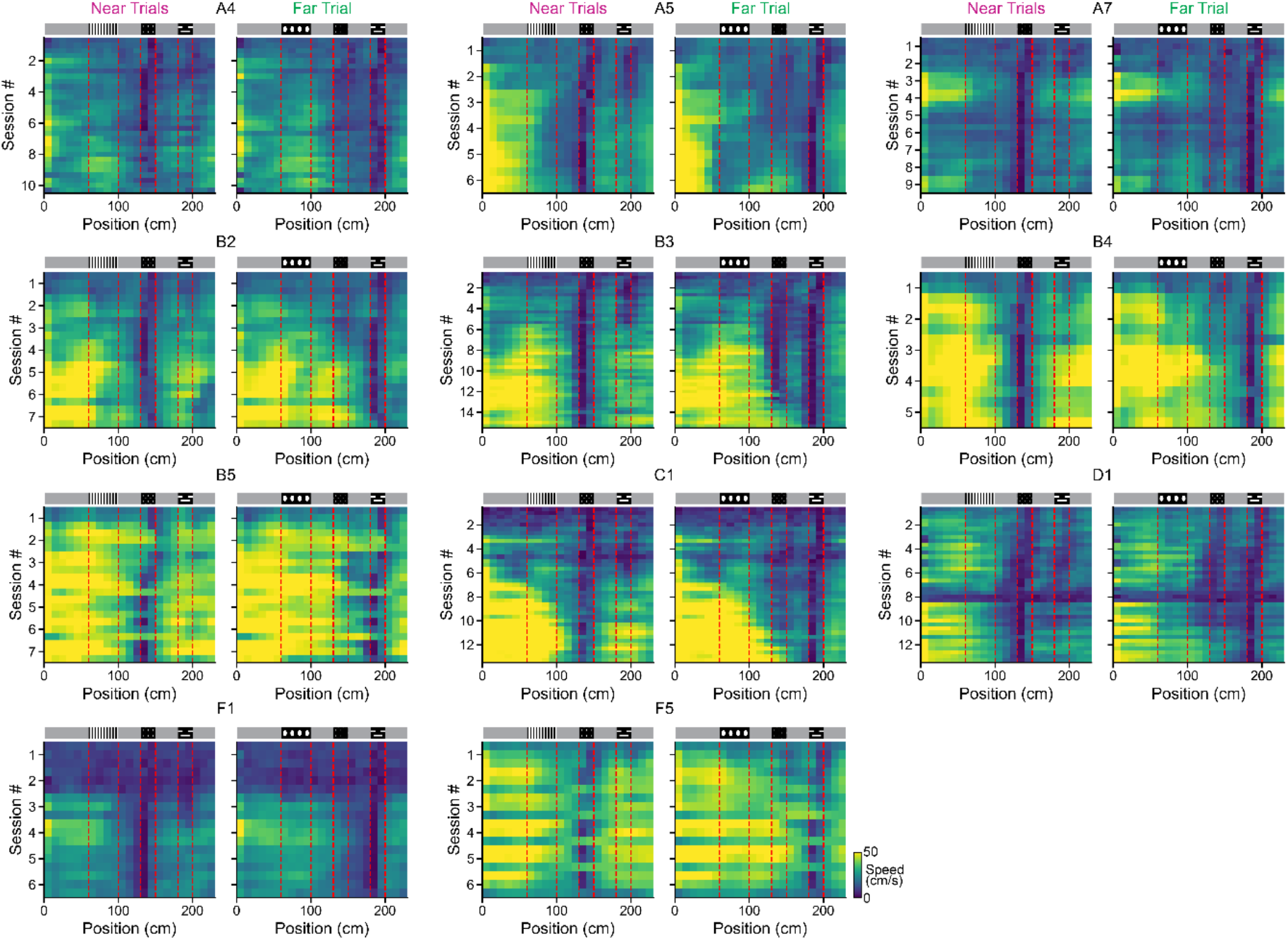
Changes in running speed during learning across all animals. Spatially binned speed profiles of all animals across training sessions during ‘Near’ and ‘Far’ trial types. Dashed red lines indicate the location of the indicator and reward cues. ‘A4’, ‘A5’… denote animal nicknames.

**Supplementary figure 3.**
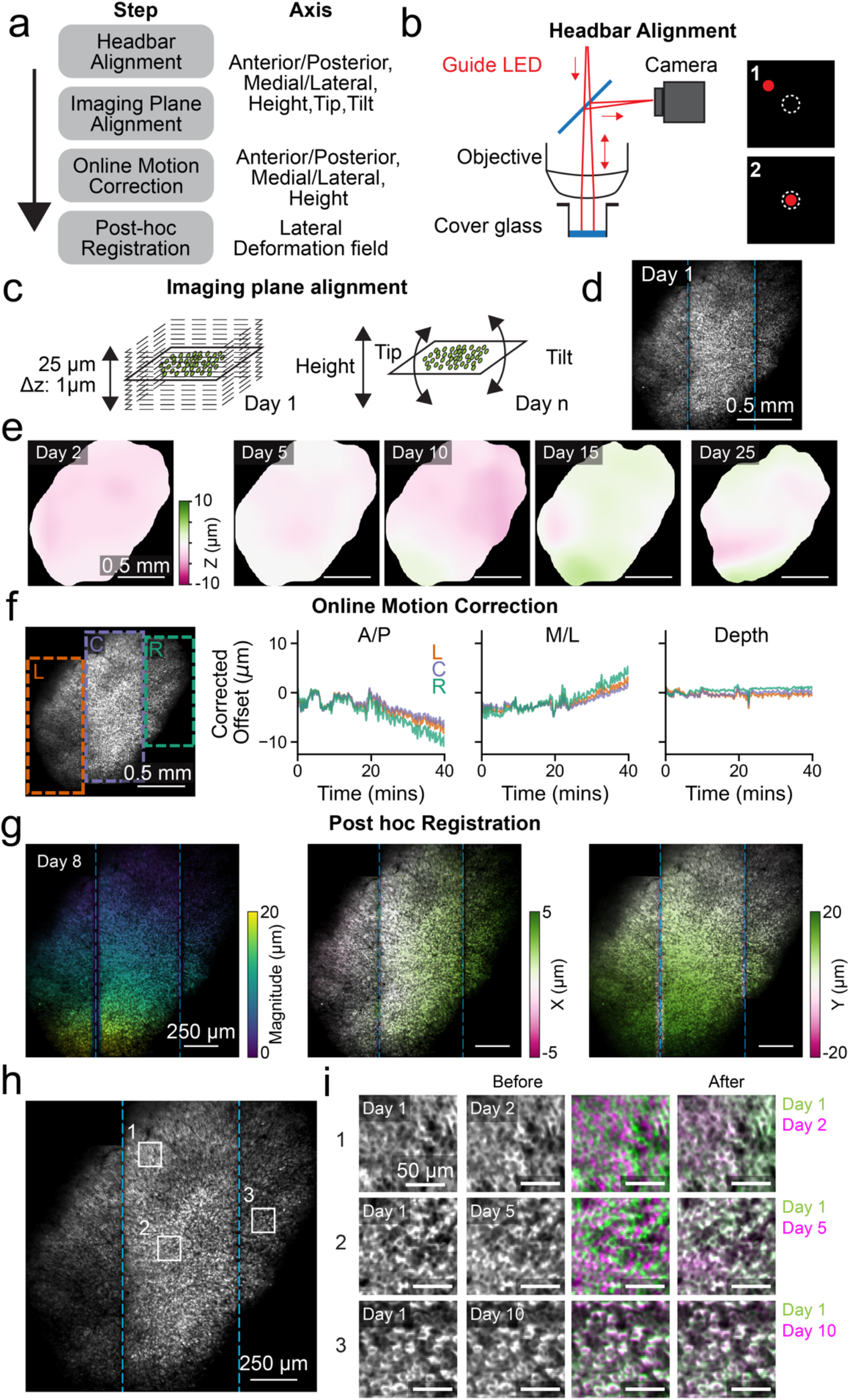
Overview of imaging alignment across sessions. (a) Left, schematic view of the alignment steps for registering the field-of-view across imaging sessions. Right, the axis that can be controlled in each alignment step. (b) Headbar alignment, focused light from a guide LED was projected through the optical path and reflected of the cover glass in the implanted canula. The position of the resulting spot on the camera sensor was used to ensure consistent alignment relative to the cover glass. (c) Image plane alignment. (left) A reference image z-stack was taken on day 1 of training. (right) The heigh, tip, and tilt of the imaging surface was adjusted on each day to achieve optimal alignment to the reference stack. (d) Example image of the field-of-view on day 1 from same animal shown in e. Dashed lines indicate the location of the imaging stripes (see Methods). (e) Heatmap of the remaining z error after alignment. (f) Online motion correction. (left) Locations of the left (orange), center (purple), and right (green) imaging stripes. (right) Online adjustment of individual imaging stripe positions during an example recording. (g) Example of post hoc registration. (left) Magnitude of elastic, non-rigid, deformation across the field-of-view. (right) Amount of deformation in x and y. (h) Location of ROIs shown in i. (i) Result of post hoc registration step. Each row shows a single ROI comparing the image on day 1 to that of day 2,5, or 10. Third and fourth column shows the overlay of the two images with day 1 in green and the comparison day in magenta both before and after registration. Note that the outline of the cells now overlap (white pixels) indicating that the same cells can be monitored across days.

**Supplementary figure 4.**
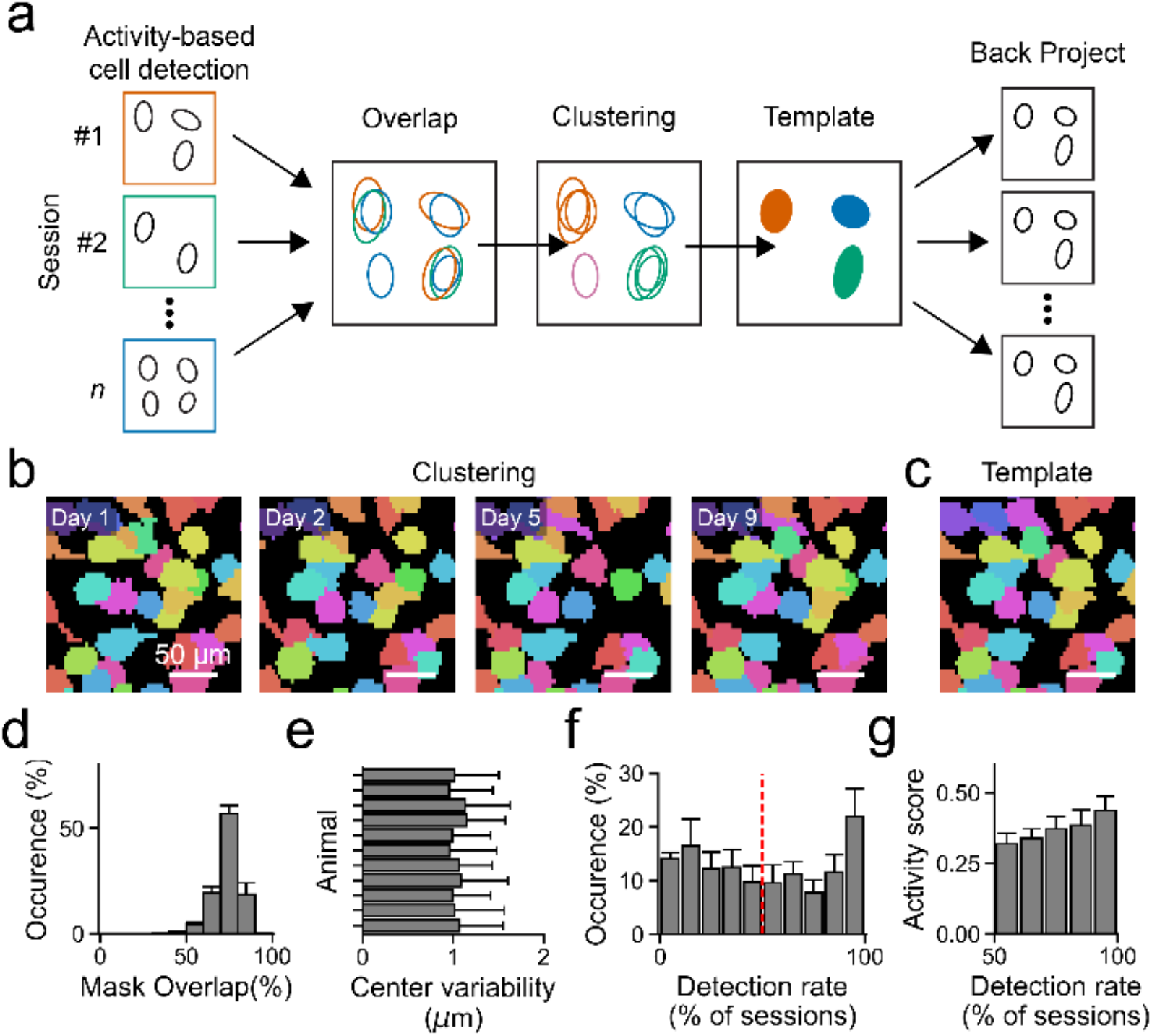
Identification of consistent cell masks across imaging sessions. (a) Schematic of computational pipeline. (left) Activity-based cell mask extraction is performed for each individual session using Suite2p (see methods). (center) The overlap between identified cell masks across all imaging sessions after registration was calculated and used to perform hierarchical clustering. The resulting clusters are used to calculate a single ‘template’ cell mask based on the median of present pixels across all sessions. Cell mask clusters that were not detected in the majority of sessions, or whose template mask was too small, were discarded (see methods for additional details). (right) The template masks were projected back to the spatial reference frame of each individual imaging session and used for calcium trace extraction. (b) Example of clustered cell masks across imaging sessions. (c) Resulting cell mask templates of the same cells shown in b. (d) Histogram of the percentage of spatial overlap of clustered cell masks with their resulting template mask averaged across animal€(e) Average variability in the center of clustered cell masks for all animals. (f) Histogram of observed detection rates of clustered cell masks as percentage of all sessions averaged across animals. Red line indicates the used inclusion threshold detection rate (50%) for all further analysis. (g) Relationship between cell detection rate and a cells activity score determined as the averaged deconvolved fluorescence signal across all sessions. Histogram values are averaged across all animals.

**Supplementary figure 5.**
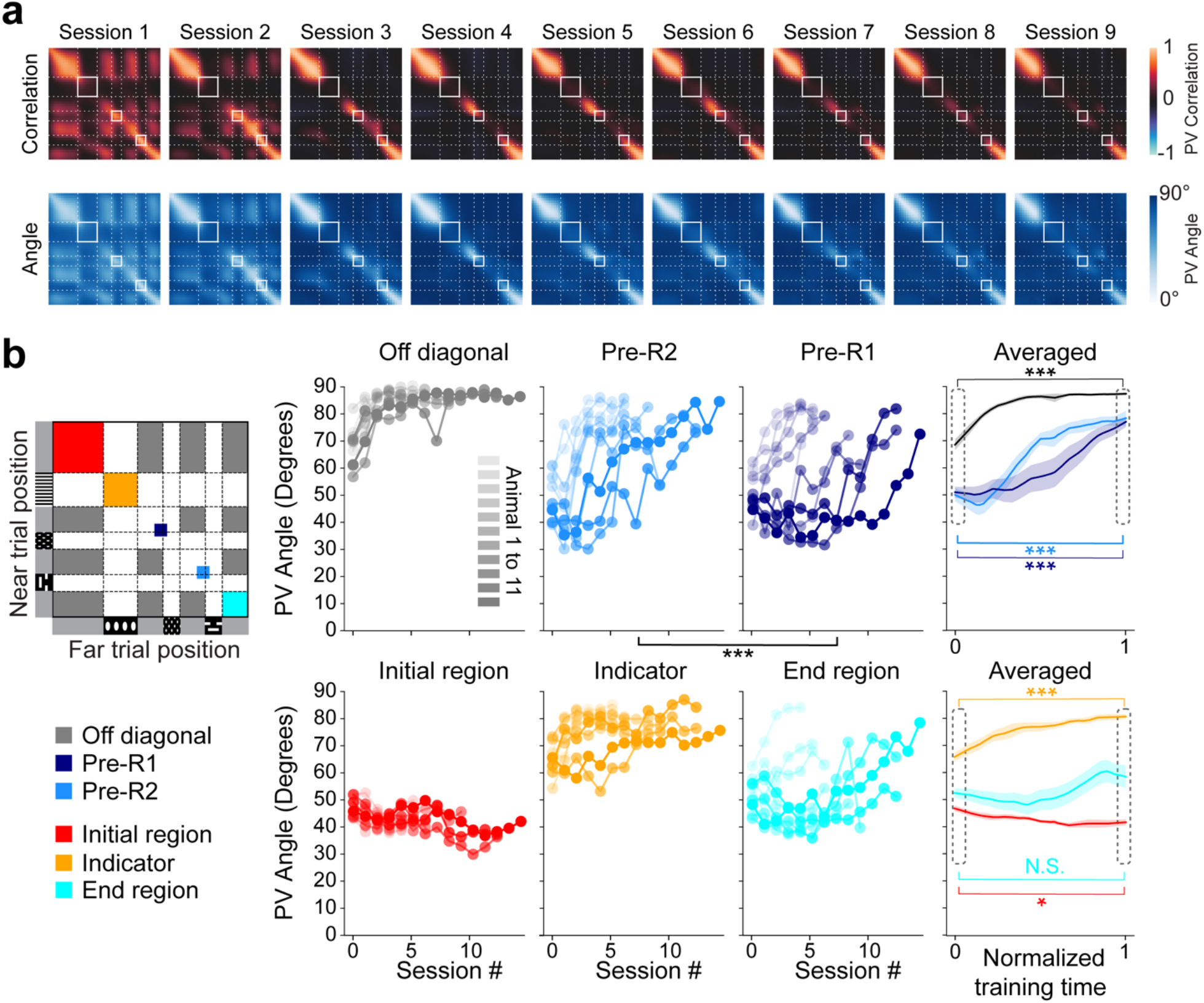
Changes in PV angle mirror the progressive decorrelation dynamics. (a) Top: *Near*-versus-*Far* PV Cross-correlation matrices along all track positions for sessions 1, 2, 3, 4, 9 for an example animal. Bottom: *Near*-versus-*Far* PV angle matrices along all track positions for the same animal. (b) PV angle averaged for different regions on the angle matrix across sessions, for the off-diagonal gray region correlations (gray), Pre-R2 region (light blue), and Pre-R1 region (dark blue), Initial region (red), Indicator region (orange), End region (cyan) shown for each animal separately and averaged across all of them. Comparing all sessions, a significant difference was observed between the Pre-R1 and Pre-R2 regions (Wilcoxon signed-rank test, *P* < 0.001***). Comparisons between the first and last session revealed significant changes in Off-diagonal gray regions, Pre-R2, Pre-R1, Indicator (angle increasing over sessions), and Initial region (angle decreasing over sessions) with *P* < 0.05*. In contrast, changes in the End region were not significant (N.S.). These results qualitatively mirror those in Fig. 2f.

**Supplementary figure 6.**
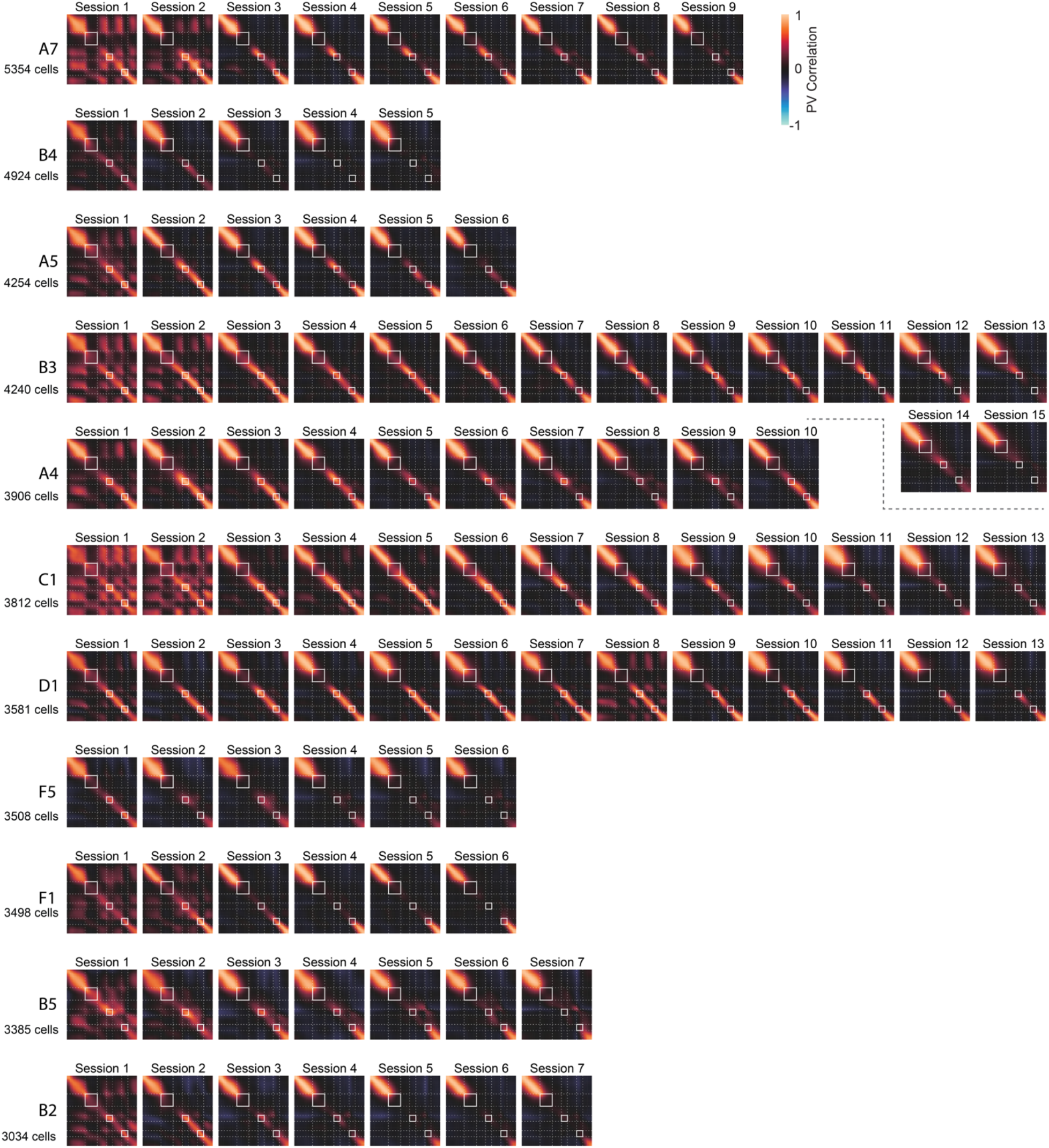
*Near* vs *Far* trial cross-correlation matrices for all 11 animals through all training sessions. Animals are ordered by the number of cells registered across sessions.

**Supplementary figure 7.**
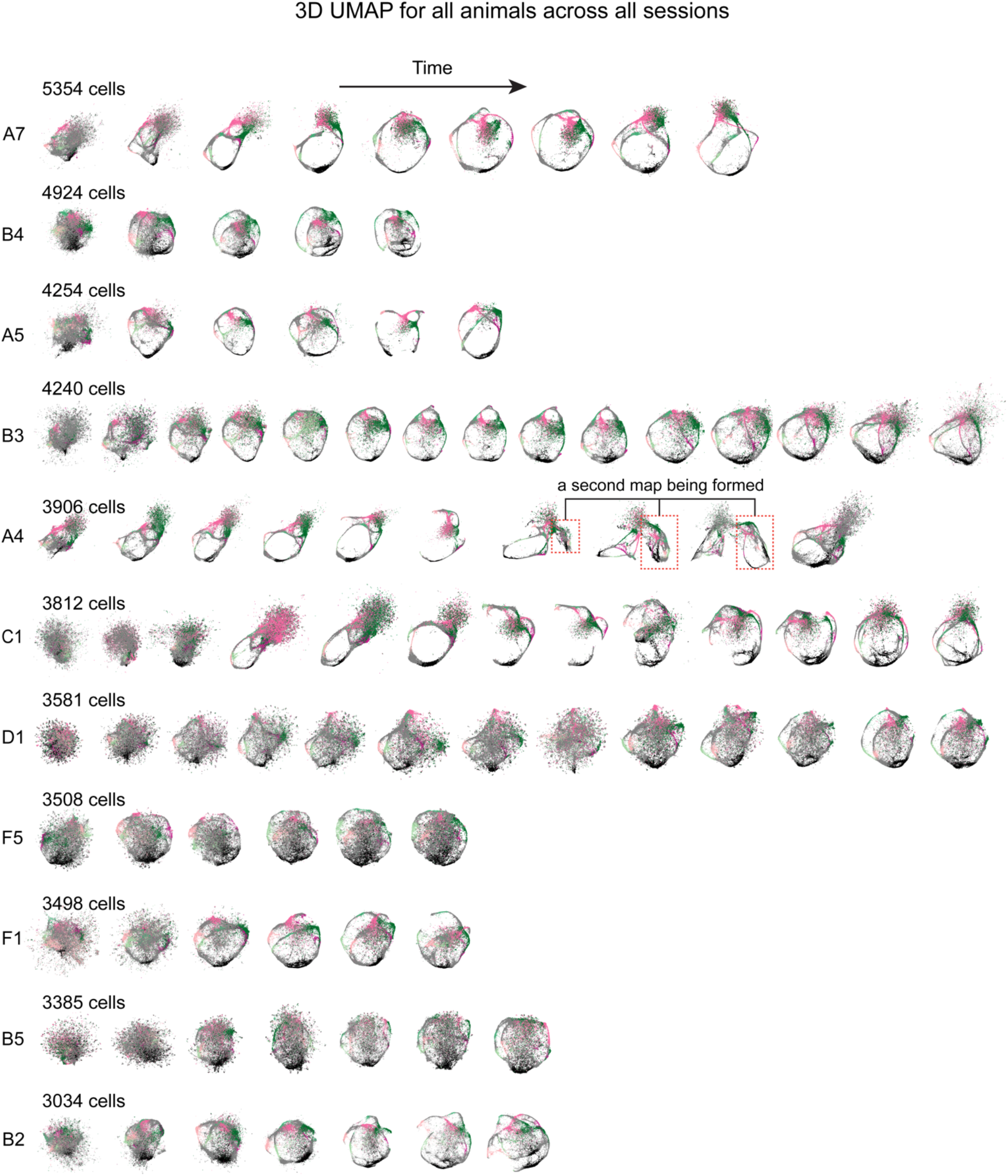
UMAP for all 11 animals through all training sessions. Animals are ordered by the number of cells registered across sessions. Note that while UMAPs shed light on the dynamics of neural activity, our conclusions are primarily driven by the representational structure reflected by the PV angles and PV correlations (Suppl. Fig. 5, 6). The utility of UMAP, influenced by the choice of hyperparameters and cell count, can yield a range of representations. Some may appear visually streamlined while others might seem noisy or fragmented. Even though their visual presentation may differ, these manifolds can offer potential insights into underlying neural dynamics. For example, the discovered manifolds can help reveal individual variability. In some animals, UMAP and correlation matrices both indicated lack of decorrelation at the track’s end (Suppl. Fig. 6). In other cases, UMAP revealed otherwise less visible aspects, such as error trials showing single trial UMAP trajectory jumping between the embeddings of correct *Near* and *Far* trial types and a novel map appearing during learning (animal A4, this form of ‘remapping’ in an unchanging environment has also been observed and modeled in the entorhinal cortex^143,144^).

**Supplementary figure 8.**
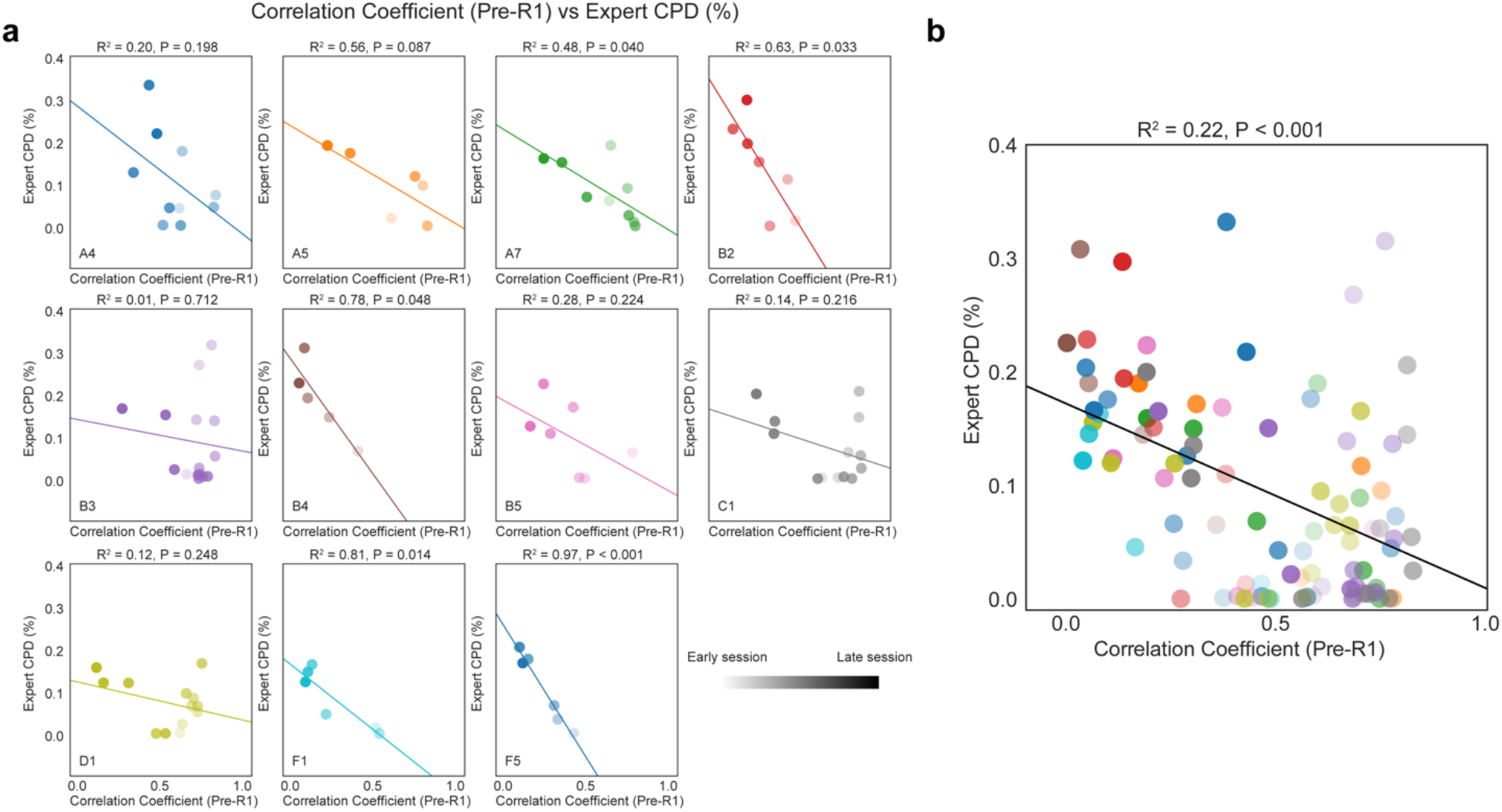
Licking behavior and neural decorrelation coevolve during learning. (a) Correlation coefficient for the Pre-R1 region between the *Near* and *Far* trials plotted against the CPD (%) for the ‘expert’ basis function for all sessions for each animal. The transparency of the filled dots indicates stage of training, with earlier sessions more transparent. Lines indicate linear regression fits, with R^2^ and P values shown on top of each plot. (b) Same with (a), but with pooled data from all animals.

**Supplementary figure 9.**
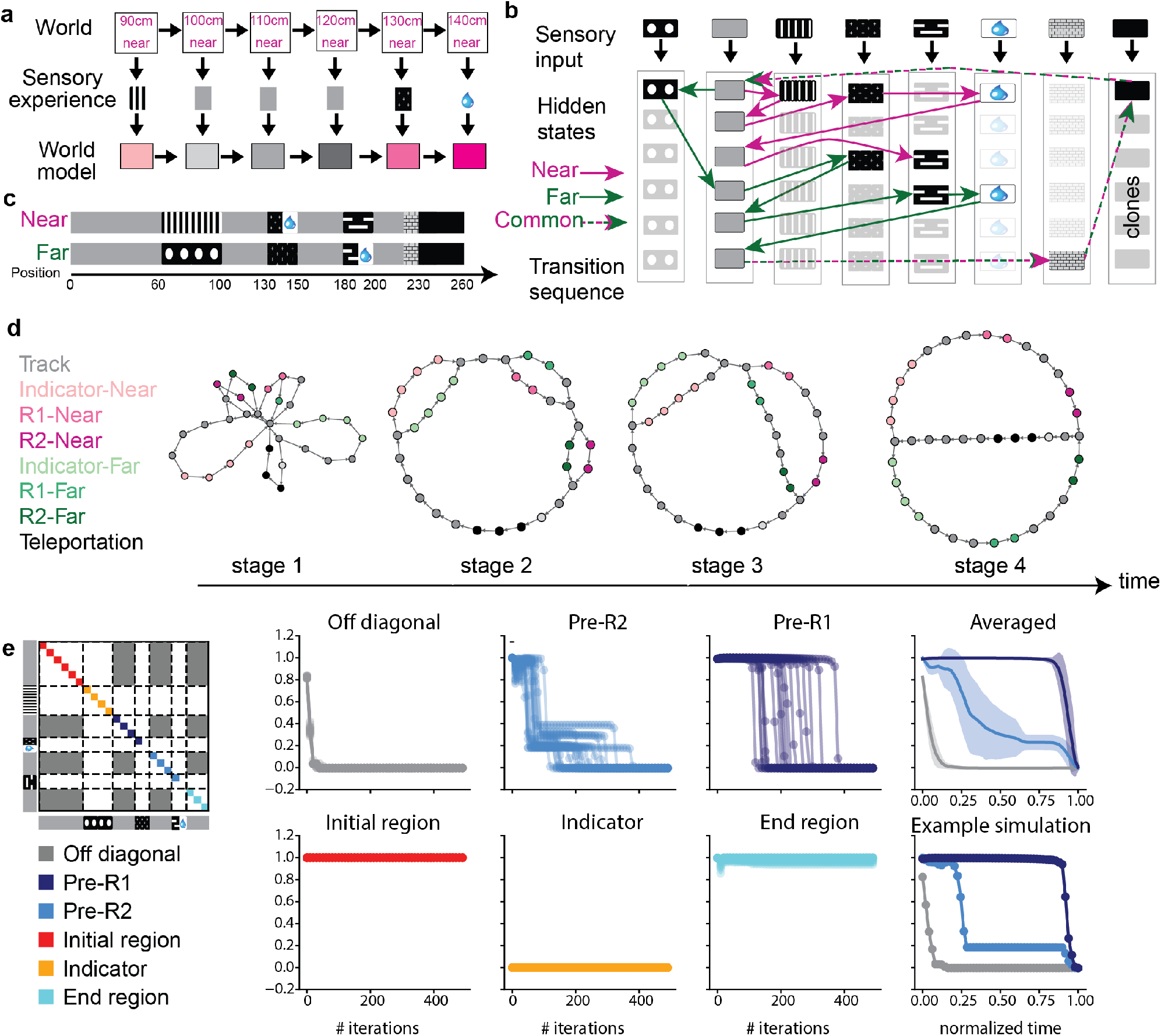
Hidden Markov learning using Clone-Structured Causal Graph recapitulates animal’s learning process. (a) The world state, determined by the position and trial type, is not directly accessible to the model. Instead, the system can access sensory experiences generated based on the world state, which is used to learn a world model that accurately predicts the next sensory experience. (b) Schematic of the CSCG and the learned transition sequence. Each sensory stimulus is associated with a set of clones, which determine the hidden state. The system learns transition probabilities between these clones to generate a world model. Gray sensory stimuli are observed at distinct locations on the near and far trials, and different clones representing the gray stimulus learn to represent distinct locations. For less ambiguous stimuli, such as the indicator, most clones remain unused. (c) Schematic representation of the sequence of sensory stimuli presented to the model for near and far trial types. The rewarded region R1 of near trial and R2 of far trial are divided into an initial visual symbol common to both trial types, followed by a shared reward symbol. The end of track symbol simulates the wall observed by the animal before teleportation. (d) The transition graph of CSCG during different learning stages recapitulates the low-dimensional neural manifolds observed in animals during learning. (e) Matrix depicting the correlation of probabilities over clones averaged for different regions over time during learning. The off-diagonal gray region (gray) decorrelates first, followed by the Pre-R2 region (sky blue), which decorrelates to a non-zero value, and then when the Pre-R1 region (navy blue) decorrelates, it is accompanied by complete decorrelation of the Pre-R2 region. The initial and end regions remain correlated. Simulations that did not fully decorrelate both Pre-R1 and Pre-R2 were excluded.

**Supplementary figure 10.**
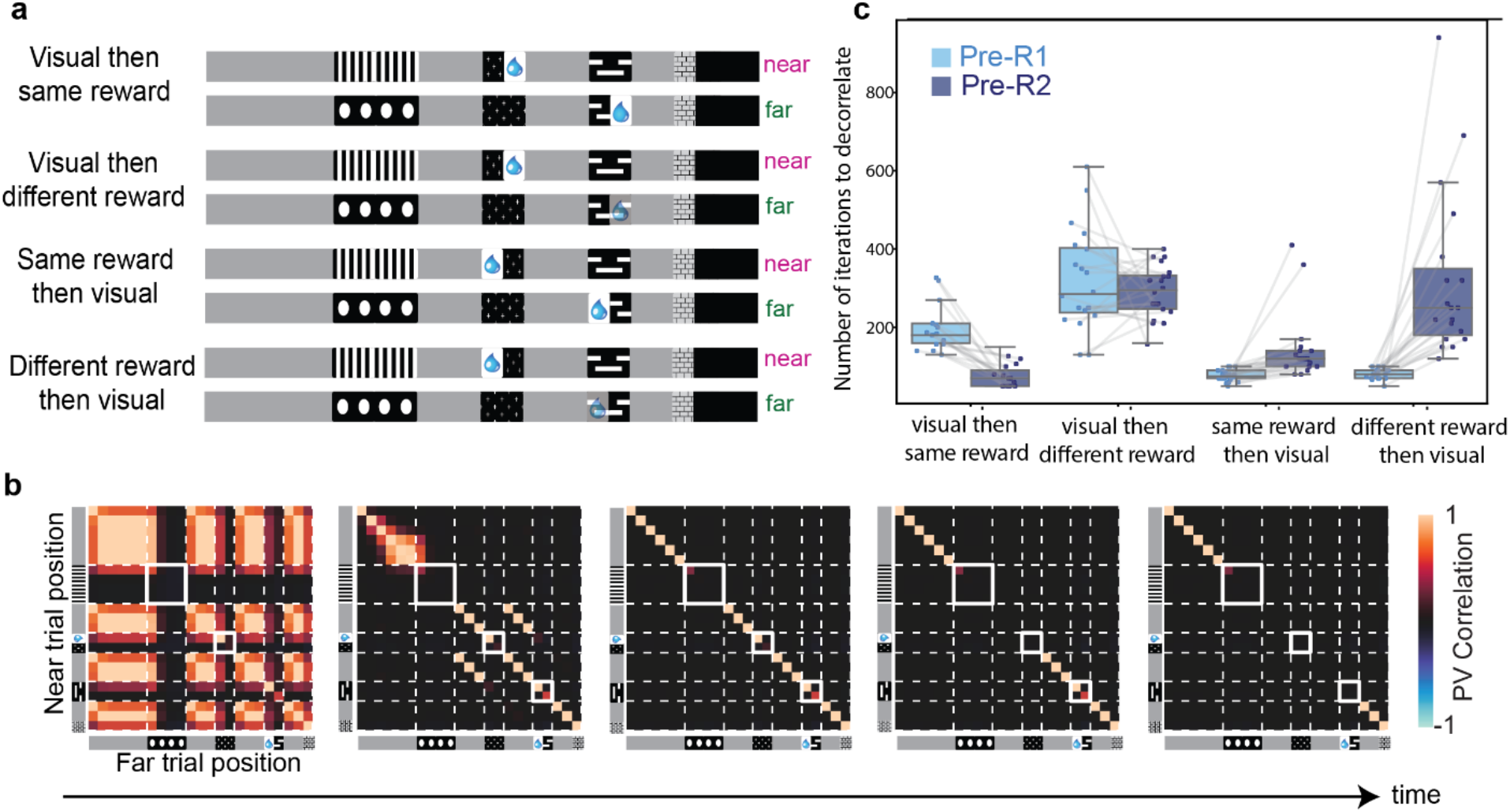
Sequence of learning/decorrelation depends on the exact sequence of sensory stimuli used in training CSCG. (a) Schematic representation of different possible sensory symbol sequences mimicking the animal’s experience, including different orders of visual and reward experiences, and a separate reward or a combined code for reward and visual symbol. (b) Cross-correlation matrices illustrating the near-versus-far probability over clones along all track positions for the reward followed by visual sequence, showing a different learning trajectory with Pre-R1 decorrelating first, followed by Pre-R2. (c) Time taken for the correlation between vectors of probability over clones (of Pre-R1 and Pre-R2 regions) between the near and far trial types to drop below 0.5. An asymmetry between Pre-R1 and Pre-R2 regions is created when the reward is followed by a visual symbol, as the system needs to decorrelate 1 R1 visual and 3 gray zones Pre-R2, whereas only 3 gray zones Pre-R1 need to be decorrelated, thus leading to decorrelation of Pre-R1 first. A visual symbol followed by reward suggests an equal probability of either Pre-R1 or Pre-R2 decorrelation occurring first, but a shared reward symbol makes it more likely for Pre-R2 to decorrelate first, mimicking animal’s learning trajectory.

**Supplementary figure 11.**
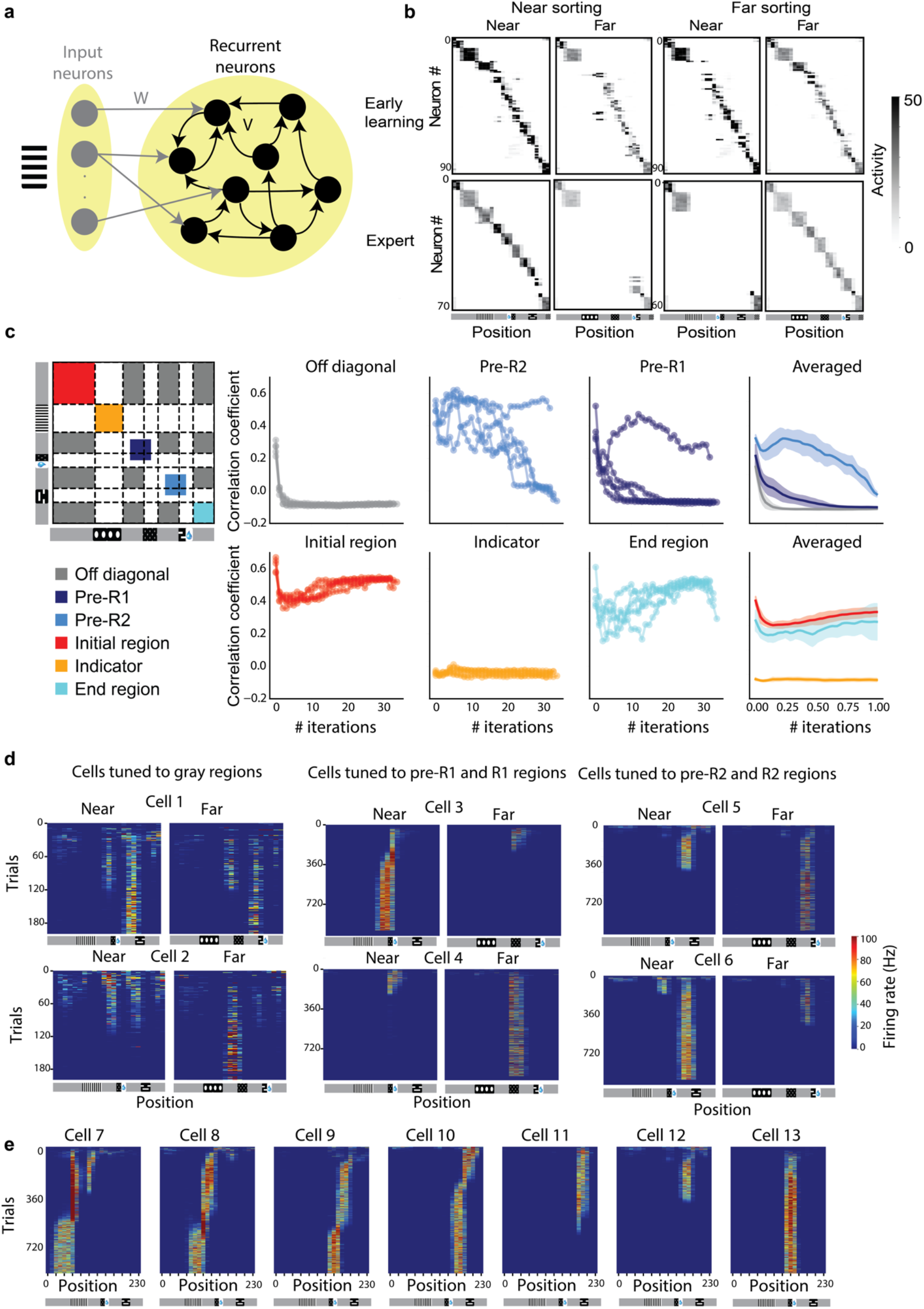
Hebbian-RNN recapitulates learned representations of animals at the population and single-cell level, though the precise learning trajectory differs between animals and RNN. (a) Schematic representation of a recurrent neural network (RNN) used to model the hippocampus. (b) Trial-averaged neural activity plotted against track position for both near and far trial types, at early and late stages of learning. Left: Cells ordered by their activity in the near trial type. Right: Cells ordered by their activity in the far trial type. Initially, the same cells encode both trial types (except the indicator region), but as learning progresses, cells coding for regions from the indicator to R2 become trial type specific. (c) Near vs far PV (Population Vector) Matrix depicting the correlation of probabilities over clones averaged for different regions over time during learning. The off-diagonal gray region (gray) decorrelates first, followed by the pre-R1 region (navy blue), and then pre-R2 region (sky blue). This suggests an order different from that observed in most animals. The initial and end regions show a slight decrease and subsequent slight increase in correlation, while the indicator region remains uncorrelated. (d) Dynamics of positional tuning for RNN cells replicate aspects of the single-cell dynamics observed in animals. Left: Example cells involved in the transition from stage 1 to stage 2, where neurons tuned to multiple gray regions become selective to one. Middle: Example cells tuned to pre-R1 and R1 regions for both trial types become selective to one trial type. Right: Example cells tuned to pre-R2 and R2 regions for both trial types become selective to one trial type. (e) Example cells exhibiting selective firing at various locations along the track in the near trial type. This includes a backward shift in cells 7 to 10, loss of selectivity in cells 11 and 12, and a stable field in cell 13.

**Supplementary figure 12.**
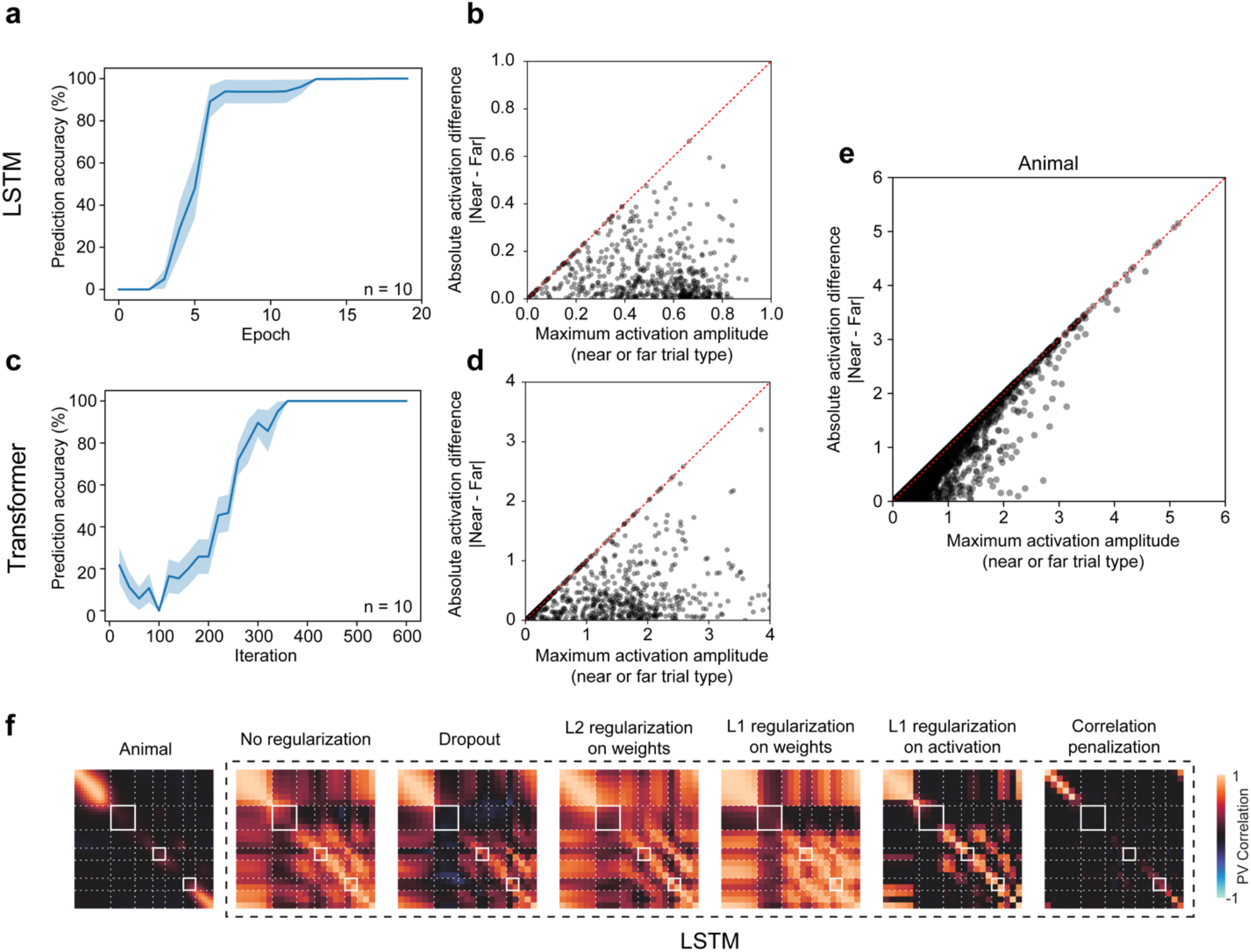
Training LSTM and Transformer on 2ACDC task. Prediction accuracy for reward regions in both *Near* and *Far* trials for LSTM (a), and Transformer (c), both shown average of 10 independent runs, shading indicates ± s.e.m. Prediction accuracy is defined as the percentage of correct reward region predictions (for both *Far* and *Near* trial types) out of all reward region predictions during test trials. (b) Maximum unit activation in *Near* and *Far* trials at a particular location plotted against the absolute activation difference between *Near* and *Far* trials, for all units and pooled across all positions from the beginning of R1 to end of R2. (d) Similar in b, but for transformer pre-logit layer activations. (e) Similar to b, but for animal neuronal activations. (f) Final model representational structure using different regularizations applied to LSTM. Regularization strength was incremented progressively; the final level was selected when subsequent increase began to degrade test performance. Correlation penalization involved storing hidden state activations for both a *Near* and a *Far* trial. The sum of all entries within the cross-correlation matrix between the two trial types was then added to the training loss.

**Supplementary figure 13.**
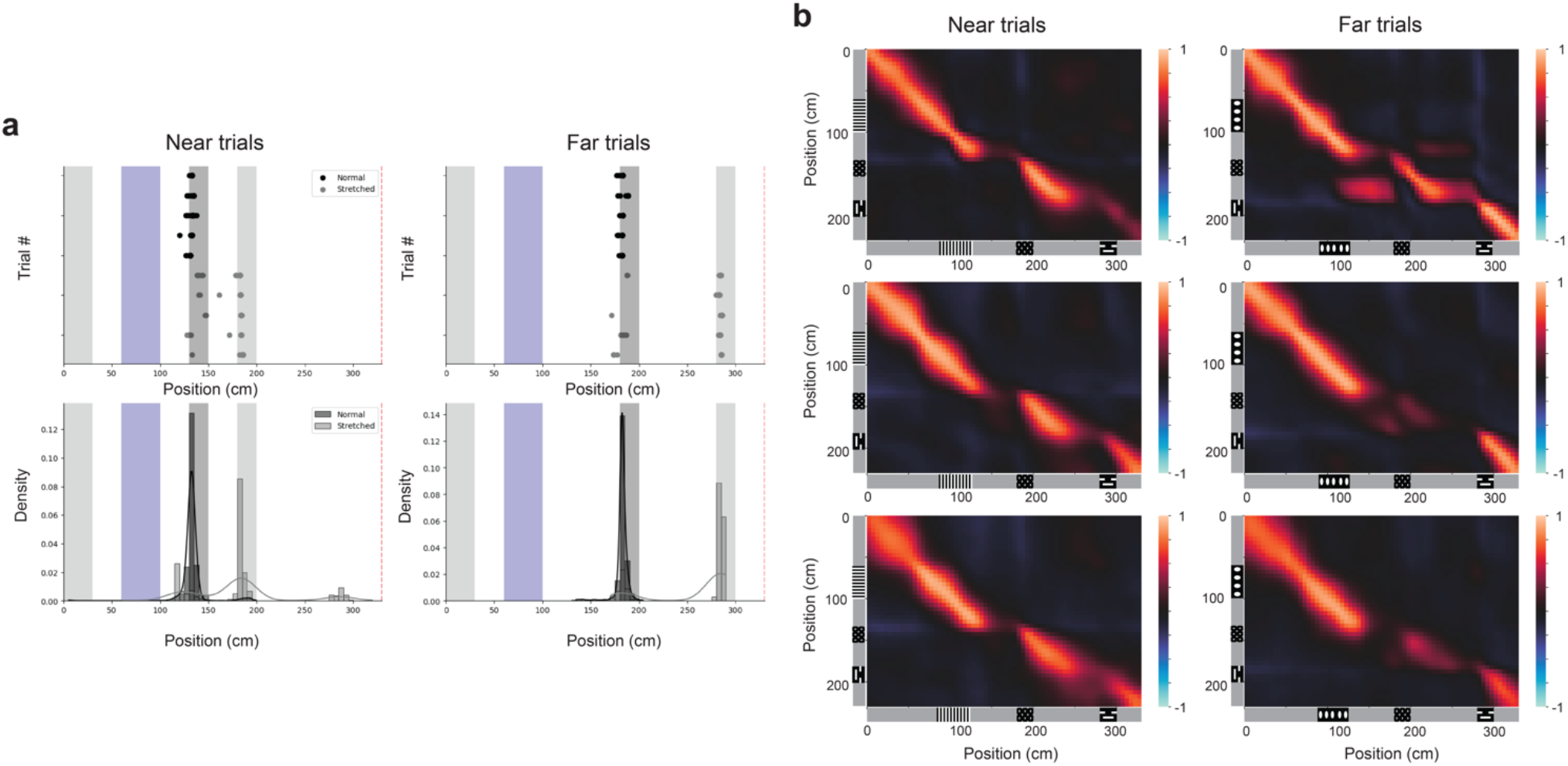
Behavior and neural activity in stretched trials. (a) Example licking patterns (top row) and the licking position distribution over a single session (bottom row) in both near and far trials for normal (black) and stretched (gray) trials. (b) PV Correlation between the average neural population activity in normal and in stretched trials for both near (left column) and far (right column) trials. Each row corresponds to a single animal.

## Notes

### Competing Interest Statement

The authors have declared no competing interest.

### Summary of Updates

Updates in main text, figures, and author list.

## REFERENCES

1. Tolman, E. C. Cognitive maps in rats and men. Psychol. Rev. 55, 189–208 (1948).

2. O’Keefe, J. & Dostrovsky, J. The hippocampus as a spatial map. Preliminary evidence from unit activity in the freely-moving rat. Brain Res. 34, 171–175 (1971).

3. O’Keefe, J. Place units in the hippocampus of the freely moving rat. Exp. Neurol. 51, 78–109 (1976).

4. O’Keefe, J. & Nadel, L. The Hippocampus as a Cognitive Map. (1978).

5. Moser, M.-B., Rowland, D. C. & Moser, E. I. Place Cells, Grid Cells, and Memory. Cold Spring Harb. Perspect. Biol. 7, a021808 (2015).

6. Nielson, D. M., Smith, T. A., Sreekumar, V., Dennis, S. & Sederberg, P. B. Human hippocampus represents space and time during retrieval of real-world memories. Proc. Natl. Acad. Sci. U. S. A. 112, 11078–11083 (2015).

7. Tavares, R. M. et al. A Map for Social Navigation in the Human Brain. Neuron 87, 231–243 (2015).

8. Hsieh, L.-T., Gruber, M. J., Jenkins, L. J. & Ranganath, C. Hippocampal activity patterns carry information about objects in temporal context. Neuron 81, 1165–1178 (2014).

9. Ekstrom, A. D. et al. Cellular networks underlying human spatial navigation. Nature 425, 184–188 (2003).

10. Miller, J. F. et al. Neural Activity in Human Hippocampal Formation Reveals the Spatial Context of Retrieved Memories. Science 342, 1111–1114 (2013).

11. Quiroga, R. Q., Reddy, L., Kreiman, G., Koch, C. & Fried, I. Invariant visual representation by single neurons in the human brain. Nature 435, 1102–1107 (2005).

12. Bausch, M. et al. Concept neurons in the human medial temporal lobe flexibly represent abstract relations between concepts. Nat. Commun. 12, 6164 (2021).

13. Gulli, R. A. et al. Context-dependent representations of objects and space in the primate hippocampus during virtual navigation. Nat. Neurosci. 23, 103–112 (2020).

14. E. T. Rolls & J. -Z. Xiang. Spatial View Cells in the Primate Hippocampus and Memory Recall. Rev. Neurosci. 17, 175–200 (2006).

15. Baraduc, P., Duhamel, J.-R. & Wirth, S. Schema cells in the macaque hippocampus. Science 363, 635–639 (2019).

16. Wirth, S., Baraduc, P., Planté, A., Pinède, S. & Duhamel, J.-R. Gaze-informed, task-situated representation of space in primate hippocampus during virtual navigation. PLOS Biol. 15, e2001045 (2017).

17. Courellis, H. S. et al. Spatial encoding in primate hippocampus during free navigation. PLOS Biol. 17, e3000546 (2019).

18. Ulanovsky, N. & Moss, C. F. Hippocampal cellular and network activity in freely moving echolocating bats. Nat. Neurosci. 10, 224–233 (2007).

19. Payne, H. L., Lynch, G. F. & Aronov, D. Neural representations of space in the hippocampus of a food-caching bird. Science 373, 343–348 (2021).

20. McNaughton, B. L., Barnes, C. A. & O’Keefe, J. The contributions of position, direction, and velocity to single unit activity in the hippocampus of freely-moving rats. Exp. Brain Res. 52, 41–49 (1983).

21. Leutgeb, S., Ragozzino, K. E. & Mizumori, S. J. Convergence of head direction and place information in the CA1 region of hippocampus. Neuroscience 100, 11–19 (2000).

22. Wood, E. R., Dudchenko, P. A., Robitsek, R. J. & Eichenbaum, H. Hippocampal Neurons Encode Information about Different Types of Memory Episodes Occurring in the Same Location. Neuron 27, 623–633 (2000).

23. Frank, L. M., Brown, E. N. & Wilson, M. Trajectory Encoding in the Hippocampus and Entorhinal Cortex. Neuron 27, 169–178 (2000).

24. Aronov, D., Nevers, R. & Tank, D. W. Mapping of a non-spatial dimension by the hippocampal/entorhinal circuit. Nature 543, 719–722 (2017).

25. Nieh, E. H. et al. Geometry of abstract learned knowledge in the hippocampus. Nature 595, 80–84 (2021).

26. Dusek, J. A. & Eichenbaum, H. The hippocampus and memory for orderly stimulus relations. Proc. Natl. Acad. Sci. U. S. A. 94, 7109–7114 (1997).

27. Garvert, M. M., Dolan, R. J. & Behrens, T. E. A map of abstract relational knowledge in the human hippocampal–entorhinal cortex. eLife 6, e17086 (2017).

28. Mok, R. M. & Love, B. C. A non-spatial account of place and grid cells based on clustering models of concept learning. Nat. Commun. 10, 5685 (2019).

29. Sun, C., Yang, W., Martin, J. & Tonegawa, S. Hippocampal neurons represent events as transferable units of experience. Nat. Neurosci. 23, 651–663 (2020).

30. O’Keefe, J. & Krupic, J. Do hippocampal pyramidal cells respond to nonspatial stimuli? Physiol. Rev. 101, 1427–1456 (2021).

31. Knudsen, E. B. & Wallis, J. D. Hippocampal neurons construct a map of an abstract value space. Cell 184, 4640–4650.e10 (2021).

32. Oliva, A. Neuronal ensemble dynamics in social memory. Curr. Opin. Neurobiol. 78, 102654 (2023).

33. Stachenfeld, K. L., Botvinick, M. M. & Gershman, S. J. The hippocampus as a predictive map. Nat. Neurosci. 20, 1643–1653 (2017).

34. Whittington, J. C. R. et al. The Tolman-Eichenbaum Machine: Unifying Space and Relational Memory through Generalization in the Hippocampal Formation. Cell 183, 1249–1263.e23 (2020).

35. George, D. et al. Clone-structured graph representations enable flexible learning and vicarious evaluation of cognitive maps. Nat. Commun. 12, 2392 (2021).

36. Benna, M. K. & Fusi, S. Place cells may simply be memory cells: Memory compression leads to spatial tuning and history dependence. Proc. Natl. Acad. Sci. 118, e2018422118 (2021).

37. Recanatesi, S. et al. Predictive learning as a network mechanism for extracting low-dimensional latent space representations. Nat. Commun. 12, 1417 (2021).

38. Wilson, M. A. & McNaughton, B. L. Dynamics of the Hippocampal Ensemble Code for Space. Science (1993) doi:10.1126/science.8351520.

39. Hill, A. J. First occurrence of hippocampal spatial firing in a new environment. Exp. Neurol. 62, 282–297 (1978).

40. Frank, L. M., Stanley, G. B. & Brown, E. N. Hippocampal plasticity across multiple days of exposure to novel environments. J. Neurosci. Off. J. Soc. Neurosci. 24, 7681–7689 (2004).

41. Raju, R. V., Guntupalli, J. S., Zhou, G., Lázaro-Gredilla, M. & George, D. Space is a latent sequence: Structured sequence learning as a unified theory of representation in the hippocampus. Preprint at http://arxiv.org/abs/2212.01508 (2022).

42. Hebb, D. O. The organization of behavior: A neuropsychological theory. New York: John Wiley and Sons, Inc., 1949. 335 p. $4.00–1950 - Science Education - Wiley Online Library. https://onlinelibrary.wiley.com/doi/10.1002/sce.37303405110.

43. Rumelhart, D. E., Hinton, G. E. & Williams, R. J. Learning representations by back-propagating errors. Nature 323, 533–536 (1986).

44. Hochreiter, S. & Schmidhuber, J. Long Short-Term Memory. Neural Comput. 9, 1735– 1780 (1997).

45. Vaswani, A. et al. Attention Is All You Need. Preprint at http://arxiv.org/abs/1706.03762 (2017).

46. Smedslund, G., Arnulf, J. K. & Smedslund, J. Is psychological science progressing? Explained variance in PsycINFO articles during the period 1956 to 2022. Front. Psychol. 13, (2022).

47. Sofroniew, N. J., Flickinger, D., King, J. & Svoboda, K. A large field of view two-photon mesoscope with subcellular resolution for in vivo imaging. eLife 5, e14472 (2016).

48. McInnes, L., Healy, J. & Melville, J. UMAP: Uniform Manifold Approximation and Projection for Dimension Reduction. ArXiv180203426 Cs Stat (2020).

49. Ólafsdóttir, H. F., Bush, D. & Barry, C. The Role of Hippocampal Replay in Memory and Planning. Curr. Biol. 28, R37–R50 (2018).

50. Duvelle, É., Grieves, R. M. & Van Der Meer, M. A. Temporal context and latent state inference in the hippocampal splitter signal. eLife 12, e82357 (2023).

51. Kappel, D., Nessler, B. & Maass, W. STDP Installs in Winner-Take-All Circuits an Online Approximation to Hidden Markov Model Learning. PLoS Comput. Biol. 10, e1003511 (2014).

52. Wayne, G., et al. Unsupervised Predictive Memory in a Goal-Directed Agent. Preprint at 10.48550/arXiv.1803.10760 (2018).

53. Brunec, I. K. & Momennejad, I. Predictive Representations in Hippocampal and Prefrontal Hierarchies. J. Neurosci. 42, 299–312 (2022).

54. Miller, A. M. P. et al. Emergence of a predictive model in the hippocampus. Neuron 111, 1952–1965.e5 (2023).

55. Sherrill, K. R. et al. Generalization of cognitive maps across space and time. Cereb. Cortex 33, 7971–7992 (2023).

56. Vikbladh, O. M. et al. Hippocampal Contributions to Model-Based Planning and Spatial Memory. Neuron 102, 683–693.e4 (2019).

57. de Cothi, W. et al. Predictive maps in rats and humans for spatial navigation. Curr. Biol. 32, 3676–3689.e5 (2022).

58. Dempster, A. P., Laird, N. M. & Rubin, D. B. Maximum Likelihood from Incomplete Data via the EM Algorithm. J. R. Stat. Soc. Ser. B Methodol. 39, 1–38 (1977).

59. Markram, H., Lübke, J., Frotscher, M. & Sakmann, B. Regulation of synaptic efficacy by coincidence of postsynaptic APs and EPSPs. Science 275, 213–215 (1997).

60. Bi, G. & Poo, M. Synaptic Modifications in Cultured Hippocampal Neurons: Dependence on Spike Timing, Synaptic Strength, and Postsynaptic Cell Type. J. Neurosci. 18, 10464–10472 (1998).

61. Fiete, I. R., Senn, W., Wang, C. Z. H. & Hahnloser, R. H. R. Spike-Time-Dependent Plasticity and Heterosynaptic Competition Organize Networks to Produce Long Scale-Free Sequences of Neural Activity. Neuron 65, 563–576 (2010).

62. Okubo, T. S., Mackevicius, E. L., Payne, H. L., Lynch, G. F. & Fee, M. S. Growth and splitting of neural sequences in songbird vocal development. Nature 528, 352–357 (2015).

63. Lisman, J. E., Talamini, L. M. & Raffone, A. Recall of memory sequences by interaction of the dentate and CA3: A revised model of the phase precession. Neural Netw. 18, 1191– 1201 (2005).

64. Khona, M. & Fiete, I. R. Attractor and integrator networks in the brain. Nat. Rev. Neurosci. 23, 744–766 (2022).

65. Fang, C., Aronov, D., Abbott, L. & Mackevicius, E. L. Neural learning rules for generating flexible predictions and computing the successor representation. eLife 12, e80680 (2023).

66. George, T. M., de Cothi, W., Stachenfeld, K. L. & Barry, C. Rapid learning of predictive maps with STDP and theta phase precession. eLife 12, e80663 (2023).

67. Bono, J., Zannone, S., Pedrosa, V. & Clopath, C. Learning predictive cognitive maps with spiking neurons during behavior and replays. eLife 12, e80671 (2023).

68. Nessler, B., Pfeiffer, M., Buesing, L. & Maass, W. Bayesian Computation Emerges in Generic Cortical Microcircuits through Spike-Timing-Dependent Plasticity. PLoS Comput. Biol. 9, e1003037 (2013).

69. Cybenko, G. Approximation by superpositions of a sigmoidal function. Math. Control Signals Syst. 2, 303–314 (1989).

70. Gühring, I., Raslan, M. & Kutyniok, G. Expressivity of Deep Neural Networks. Preprint at 10.48550/arXiv.2007.04759 (2020).

71. Marblestone, A. H., Wayne, G. & Kording, K. P. Toward an integration of deep learning and neuroscience. Front. Comput. Neurosci. 10, 94 (2016).

72. Grienberger, C. & Magee, J. C. Entorhinal cortex directs learning-related changes in CA1 representations. Nature 611, 554–562 (2022).

73. Zheng, Y., Liu, X. L., Nishiyama, S., Ranganath, C. & O’Reilly, R. C. Correcting the hebbian mistake: Toward a fully error-driven hippocampus. PLOS Comput. Biol. 18, e1010589 (2022).

74. Payeur, A., Guerguiev, J., Zenke, F., Richards, B. A. & Naud, R. Burst-dependent synaptic plasticity can coordinate learning in hierarchical circuits. Nat. Neurosci. 24, 1010– 1019 (2021).

75. Cone, I. & Clopath, C. Latent Representations in Hippocampal Network Model Co-Evolve with Behavioral Exploration of Task Structure. 2023.04.24.538070 Preprint at 10.1101/2023.04.24.538070 (2023).

76. Bittner, K. C., Milstein, A. D., Grienberger, C., Romani, S. & Magee, J. C. Behavioral time scale synaptic plasticity underlies CA1 place fields. Science 357, 1033–1036 (2017).

77. Li, X.-G., Somogyi, P., Ylinen, A. & Buzsáki, G. The hippocampal CA3 network: An in vivo intracellular labeling study. J. Comp. Neurol. 339, 181–208 (1994).

78. Rolls, E. T. An attractor network in the hippocampus: Theory and neurophysiology. Learn. Mem. 14, 714–731 (2007).

79. Mishra, R. K., Kim, S., Guzman, S. J. & Jonas, P. Symmetric spike timing-dependent plasticity at CA3–CA3 synapses optimizes storage and recall in autoassociative networks. Nat. Commun. 7, 11552 (2016).

80. Sipser, M. Introduction to the Theory of Computation. ACM SIGACT News 27, 27–29 (1996).

81. Anderson, M. I. & Jeffery, K. J. Heterogeneous modulation of place cell firing by changes in context. J. Neurosci. Off. J. Soc. Neurosci. 23, 8827–8835 (2003).

82. Leutgeb, S. Independent Codes for Spatial and Episodic Memory in Hippocampal Neuronal Ensembles. Science 309, 619–623 (2005).

83. Muller, R. & Kubie, J. The effects of changes in the environment on the spatial firing of hippocampal complex-spike cells. J. Neurosci. 7, 1951–1968 (1987).

84. Leutgeb, S., Leutgeb, J. K., Treves, A., Moser, M.-B. & Moser, E. I. Distinct ensemble codes in hippocampal areas CA3 and CA1. Sci. N. Y. NY 305, 1295–1298 (2004).

85. Latuske, P., Kornienko, O., Kohler, L. & Allen, K. Hippocampal Remapping and Its Entorhinal Origin. Front. Behav. Neurosci. 11, (2018).

86. Lever, C., Wills, T., Cacucci, F., Burgess, N. & O’Keefe, J. Long-term plasticity in hippocampal place-cell representation of environmental geometry. Nature 416, 90–94 (2002).

87. Bulkin, D. A., Law, L. M. & Smith, D. M. Placing memories in context: Hippocampal representations promote retrieval of appropriate memories. Hippocampus 26, 958–971 (2016).

88. Zhao, X., Hsu, C.-L. & Spruston, N. Rapid synaptic plasticity contributes to a learned conjunctive code of position and choice-related information in the hippocampus. Neuron 110, 96–108.e4 (2022).

89. Duvelle, É., Grieves, R. M. & van der Meer, M. A. Temporal context and latent state inference in the hippocampal splitter signal. eLife 12, e82357 (2023).

90. Yassa, M. A. & Stark, C. E. L. Pattern separation in the hippocampus. Trends Neurosci. 34, 515–525 (2011).

91. Neunuebel, J. P. & Knierim, J. J. CA3 Retrieves Coherent Representations from Degraded Input: Direct Evidence for CA3 Pattern Completion and Dentate Gyrus Pattern Separation. Neuron 81, 416 (2014).

92. Knierim, J. J. & Neunuebel, J. P. Tracking the flow of hippocampal computation: Pattern separation, pattern completion, and attractor dynamics. Neurobiol. Learn. Mem. 129, 38–49 (2016).

93. Bakker, A., Kirwan, C. B., Miller, M. & Stark, C. E. L. Pattern Separation in the Human Hippocampal CA3 and Dentate Gyrus. Science 319, 1640–1642 (2008).

94. GoodSmith, D., Lee, H., Neunuebel, J. P., Song, H. & Knierim, J. J. Dentate Gyrus Mossy Cells Share a Role in Pattern Separation with Dentate Granule Cells and Proximal CA3 Pyramidal Cells. J. Neurosci. Off. J. Soc. Neurosci. 39, 9570–9584 (2019).

95. Cayco Gajic, N. A. & Silver, R. A. Re-evaluating circuit mechanisms underlying pattern separation. Neuron 101, 584–602 (2019).

96. Wiechert, M. T., Judkewitz, B., Riecke, H. & Friedrich, R. W. Mechanisms of pattern decorrelation by recurrent neuronal circuits. Nat. Neurosci. 13, 1003–1010 (2010).

97. Whittington, J. C. R. et al. The Tolman-Eichenbaum Machine: Unifying Space and Relational Memory through Generalization in the Hippocampal Formation. Cell 183, 1249–1263.e23 (2020).

98. Sanders, H., Wilson, M. A. & Gershman, S. J. Hippocampal remapping as hidden state inference. eLife 9, e51140 (2020).

99. Low, R. J., Lewallen, S., Aronov, D., Nevers, R. & Tank, D. W. Probing variability in a cognitive map using manifold inference from neural dynamics. BioRxiv 418939 (2018).

100. Sengupta, A. M., Tepper, M., Pehlevan, C., Genkin, A. & Chklovskii, D. B. Manifold-tiling Localized Receptive Fields are Optimal in Similarity-preserving Neural Networks. 338947 Preprint at 10.1101/338947 (2018).

101. Marr, D. Simple memory: a theory for archicortex. Philos. Trans. R. Soc. Lond. B. Biol. Sci. 262, 23–81 (1971).

102. Albus, J. S. A theory of cerebellar function. Math. Biosci. 10, 25–61 (1971).

103. Barlow, H. B. Possible Principles Underlying the Transformations of Sensory Messages. in Sensory Communication (ed. Rosenblith, W. A.) 216–234 (The MIT Press, 2012). doi:10.7551/mitpress/9780262518420.003.0013.

104. Kanerva, P. Sparse Distributed Memory. (MIT Press, 1988).

105. Rolls, E. T. & Treves, A. The relative advantages of sparse versus distributed encoding for associative neuronal networks in the brain. Netw. Comput. Neural Syst. 1, 407–421 (1990).

106. Olshausen, B. A. & Field, D. J. Emergence of simple-cell receptive field properties by learning a sparse code for natural images. Nature 381, 607–609 (1996).

107. Ahmad, S. & Scheinkman, L. How Can We Be So Dense? The Benefits of Using Highly Sparse Representations. Preprint at 10.48550/arXiv.1903.11257 (2019).

108. Schapiro, A. C., Turk-Browne, N. B., Botvinick, M. M. & Norman, K. A. Complementary learning systems within the hippocampus: a neural network modelling approach to reconciling episodic memory with statistical learning. Philos. Trans. R. Soc. B Biol. Sci. 372, 20160049 (2017).

109. Koay, S. A., Charles, A. S., Thiberge, S. Y., Brody, C. D. & Tank, D. W. Sequential and efficient neural-population coding of complex task information. Neuron 110, 328–349.e11 (2022).

110. Flesch, T., Juechems, K., Dumbalska, T., Saxe, A. & Summerfield, C. Orthogonal representations for robust context-dependent task performance in brains and neural networks. Neuron 110, 1258–1270.e11 (2022).

111. Zbontar, J., Jing, L., Misra, I., LeCun, Y. & Deny, S. Barlow Twins: Self-Supervised Learning via Redundancy Reduction. (2021).

112. Friston, K. Learning and inference in the brain. Neural Netw. Off. J. Int. Neural Netw. Soc. 16, 1325–1352 (2003).

113. Friston, K. The free-energy principle: a rough guide to the brain? Trends Cogn. Sci. 13, 293–301 (2009).

114. Rao, R. P. & Ballard, D. H. Predictive coding in the visual cortex: a functional interpretation of some extra-classical receptive-field effects. Nat. Neurosci. 2, 79–87 (1999).

115. Schapiro, A. C., Turk-Browne, N. B., Norman, K. A. & Botvinick, M. M. Statistical learning of temporal community structure in the hippocampus. Hippocampus 26, 3–8 (2016).

116. Gershman, S. J. & Niv, Y. Learning latent structure: carving nature at its joints. Curr. Opin. Neurobiol. 20, 251–256 (2010).

117. Whittington, J. C. R., McCaffary, D., Bakermans, J. J. W. & Behrens, T. E. J. How to build a cognitive map. Nat. Neurosci. 25, 1257–1272 (2022).

118. Niv, Y. Learning task-state representations. Nat. Neurosci. 22, 1544–1553 (2019).

119. Gluck, M. A. & Myers, C. E. Hippocampal mediation of stimulus representation: A computational theory. Hippocampus 3, 491–516 (1993).

120. van de Ven, G. M., Siegelmann, H. T. & Tolias, A. S. Brain-inspired replay for continual learning with artificial neural networks. Nat. Commun. 11, 4069 (2020).

121. Uria, B. et al. A model of egocentric to allocentric understanding in mammalian brains. 2020.11.11.378141 Preprint at 10.1101/2020.11.11.378141 (2022).

122. Foster, D. J. & Wilson, M. A. Reverse replay of behavioural sequences in hippocampal place cells during the awake state. Nature 440, 680–683 (2006).

123. Haga, T. & Fukai, T. Recurrent network model for learning goal-directed sequences through reverse replay. eLife 7, e34171 (2018).

124. McClelland, J. L., McNaughton, B. L. & O’Reilly, R. C. Why there are complementary learning systems in the hippocampus and neocortex: insights from the successes and failures of connectionist models of learning and memory. Psychol. Rev. 102, 419–457 (1995).

125. Kumaran, D., Hassabis, D. & McClelland, J. L. What Learning Systems do Intelligent Agents Need? Complementary Learning Systems Theory Updated. Trends Cogn. Sci. 20, 512–534 (2016).

126. Sun, W., Advani, M., Spruston, N., Saxe, A. & Fitzgerald, J. E. Organizing memories for generalization in complementary learning systems. BioRxiv 2021–10 (2021).

127. Tsodyks, M. V. Associative Memory in Asymmetric Diluted Network with Low Level of Activity. Europhys. Lett. 7, 203 (1988).

128. Ma, Y., Tsao, D. & Shum, H.-Y. On the Principles of Parsimony and Self-Consistency for the Emergence of Intelligence. Preprint at http://arxiv.org/abs/2207.04630 (2022).

129. Richards, B. A. et al. A deep learning framework for neuroscience. Nat. Neurosci. 22, 1761–1770 (2019).

130. Saxe, A., Nelli, S. & Summerfield, C. If deep learning is the answer, what is the question? Nat. Rev. Neurosci. 22, 55–67 (2021).

131. Dana, H. et al. Thy1-GCaMP6 Transgenic Mice for Neuronal Population Imaging In Vivo. PLOS ONE 9, e108697 (2014).

132. Cohen, J. D., Bolstad, M. & Lee, A. K. Experience-dependent shaping of hippocampal CA1 intracellular activity in novel and familiar environments. eLife 6, e23040 (2017).

133. Lopes, G. et al. Bonsai: an event-based framework for processing and controlling data streams. *Front*. Neuroinformatics 9, (2015).

134. Pologruto, T. A., Sabatini, B. L. & Svoboda, K. ScanImage: Flexible software for operating laser scanning microscopes. Biomed. Eng. OnLine 2, 13 (2003).

135. Stringer, C. et al. Spontaneous behaviors drive multidimensional, brainwide activity. Science 364, eaav7893 (2019).

136. Stringer, C., Michaelos, M., Tsyboulski, D., Lindo, S. E. & Pachitariu, M. High-precision coding in visual cortex. Cell 184, 2767–2778.e15 (2021).

137. Dombeck, D. A., Harvey, C. D., Tian, L., Looger, L. L. & Tank, D. W. Functional imaging of hippocampal place cells at cellular resolution during virtual navigation. Nat. Neurosci. 13, 1433–1440 (2010).

138. Grijseels, D. M., Shaw, K., Barry, C. & Hall, C. N. Choice of method of place cell classification determines the population of cells identified. PLOS Comput. Biol. 17, e1008835 (2021).

139. Baum, L. E., Petrie, T., Soules, G. & Weiss, N. A Maximization Technique Occurring in the Statistical Analysis of Probabilistic Functions of Markov Chains. Ann. Math. Stat. 41, 164–171 (1970).

140. Do, C. B. & Batzoglou, S. What is the expectation maximization algorithm? Nat. Biotechnol. 26, 897–899 (2008).

141. Ghojogh, B., Karray, F. & Crowley, M. Hidden Markov Model: Tutorial. Preprint at 10.31224/osf.io/w9v2b (2019).

142. Srivastava, N., Hinton, G., Krizhevsky, A., Sutskever, I. & Salakhutdinov, R. Dropout: A Simple Way to Prevent Neural Networks from Overfitting. J. Mach. Learn. Res. 15, 1929– 1958 (2014).

143. Low, I. I. C., Williams, A. H., Campbell, M. G., Linderman, S. W. & Giocomo, L. M. Dynamic and reversible remapping of network representations in an unchanging environment. Neuron 109, 2967–2980.e11 (2021).

144. Low, I. I., Giocomo, L. M. & Williams, A. H. Remapping in a recurrent neural network model of navigation and context inference. eLife 12, RP86943 (2023).

